# Insights into strepsipteran flight

**DOI:** 10.64898/2026.01.26.701776

**Authors:** Marisano James

**Affiliations:** University of California at Davis Department of Evolution and Ecology Davis, California, United States of America

**Keywords:** sex pheromone, wing loading, power loading, flight muscle ratio (FMR), halteres, vision

## Abstract

Because Strepsiptera can fly vertically from a standing start, at least ¼ of their body mass must be dedicated to flight muscle. Adult male Strepsiptera also do not feed and die within a few hours of eclosing, so much normal adult insect anatomy has been discarded, leading to a flight muscle to total mass ratio (FMR) of at least 30%—this is medial for Hymenoptera, but as the *lower* bound for Strepsiptera, it indicates substantial aerial ability. On account of their high FMR and low wing loading, Strepsiptera are capable of widely varied flight. Moreover, the often incongruous descriptions thereof (that they fly slowly, fly quickly, are clumsy, are graceful, etc.) are paralleled in well-established phases of sex pheromone tracking in moths. For nearly all of their brief eclosed adult lives, male Strepsiptera are airborne, for which they are well-adapted. Correspondingly, strepsipteran propagation is utterly dependent on flight. Thus, flight is the lens through which much strepsipteran ecology is clarified. Accordingly, I photographed free-flying *Triozocera texana* (nocturnal) in the field and analyzed the images. Strepsipteran wings are remarkably flaccid and potentially teneral, leading to certain flight advantages. At night, spatial acuity is especially poor in tiny insects, but halteres apparently compensate so well that even later derived diurnal Strepsiptera identify calling females chemotactilely—not visually—and shun resolution for high sensitivity. Future directions are discussed, as well as experimental techniques that are problematic when applied to Strepsiptera.

## Introduction

There are anecdotes of the elegance and apparent leisure with which Strepsiptera fly: “The little animals [*Stylops thwaitesi*] are exceedingly graceful in their flight, taking long sweeps, as if carried along by a gentle breeze…” (Thwaites 1841). But also reports of astonishing speed: “Its [*Xenos pallidus*^1^(?)] life as an active imago cannot be longer than fifteen or twenty minutes, if as long, and during this time it exhibits fiery energy, and flies so rapidly that the eye can hardly follow it.” (Hubbard 1892). And also ostensibly contradictory accounts of its poor flight ability: “[I]ts [*Stylops perkinsi*] flight is different to anything else I have ever seen; a very peculiar flight, … what I should call an uncomfortable flight, up and down, this way and that way, in fact at all angles, not keeping in one direction more than a few inches [≈ 8 cm]…” (Enock 1875). In this study, I present a reconciliation of these divergent observations.

Strepsiptera (Fig. 1) is the only insect order in which all feeding is endoparasitic.^2^ Adult females cannot fly, and except in the most basal extant lineage, cannot see, walk, or even exit their hosts. Consequently, they also cannot oviposit. The young hatch internally and consume their mother via hemocoelous viviparity (Hagan 1948). Later, the host is relied upon to inadvertently help disperse strepsipteran 1^st^ instars in areas frequented by new potential hosts. If an infected (or “stylopized”) host dies when its Strepsiptera are already pupal or adult (neither of which feed), then the parasites may survive, particularly if male (SS Saunders 1853; Dury 1902; Kirkpatrick 1937). However, diurnal adult males will only emerge if exposed to light (SS Saunders 1853).^3^ Also, a female that is able to call (i.e., release sex pheromone) from within a dead host might still mate (Kirkpatrick 1937), but would desiccate before her offspring could mature. Planidia of a parturient female embedded in a dead host can eclose if the body remains moist (Kirkpatrick 1937), but can travel less than a foot unaided (Dury 1902; Kirkpatrick 1937). Thus, if a host dies prematurely, it must do so in a serendipitous location.

**Fig. 1.**
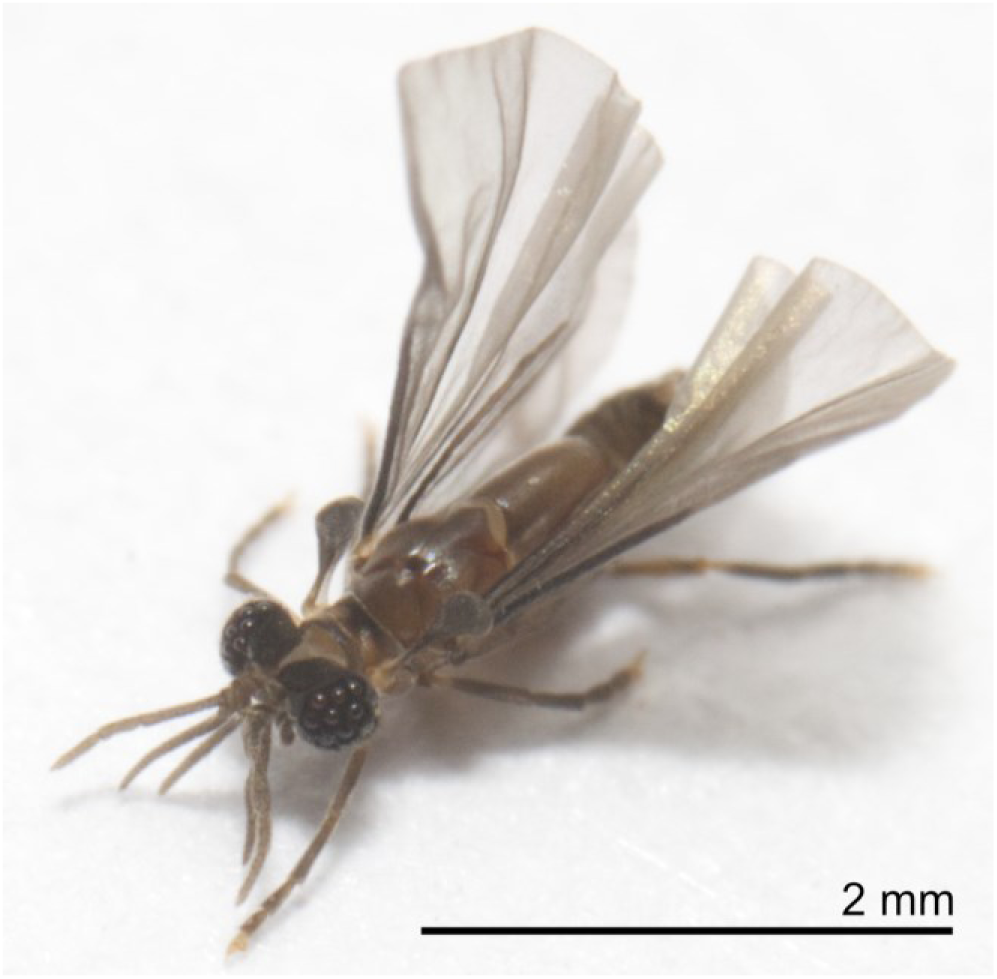
Moribund adult male *Triozocera texana*. Jumbled legs and drooped antennae indicate death is imminent. Although pharate^4^ adult males survive much longer, upon emergence, they die within about 4 h. Like true flies, Strepsiptera also have halteres (flap-mediated gyroscopic sensors) but unlike Diptera, whose halteres are modified hindwings, strepsipteran halteres are modified forewings, reflecting their phylogenetic affinity with beetles (Coleoptera). Like flies, however, airborne Strepsiptera are agile and also capable of hovering. Male *T. texana* are 2–3 mm long. Both sexes parasitize the burrowing bug *Pangaeus bilineatus*, but as with most other strepsipteran taxa, adult female *T. texana* have no wings, eyes, antennae, or legs, so after finding a host as a larva, they never leave. How their minute larvae—which cannot dig—arrive at larval hosts that live underground is unknown, but must rely on maternal care, which is well-documented in Cydnidae, to which *P. bilineatus* belongs

Thus, unlike parasitoids, which are free to kill the host in the course of their own development, and often benefit from doing so, “modern”^5^ Strepsiptera have become true parasites. Virgin females call chemically, their sex pheromone summoning conspecifics that engage in an airborne duel with dwindling time. Within a few hours, energy-depleted males may literally fall from the sky (Hubbard 1892) and twitch on the ground as they slowly die; figuratively: every parasite strepsipterous, a Siren or an Icarus. Each male is equipped to fly upon (Hrabar et al. 2014) or promptly after adult emergence (Hassan 1939; Smith & Kathirithamby 1984) using his highly protractile abdomen to push free of his puparium (Ulrich 1933), whereas most insects must pause to expand and sclerotize their wings—a process often lasting hours; time Strepsiptera do not have to spare: Their hosts move as they will, more or less burdened by their passengers, but apparently without direction or overt interference from them.

Parthenogenesis is unknown in Strepsiptera, so flight is of fundamental importance. From direct interaction with flying crepuscular *Elenchus koebelei* and nocturnal *Triozocera texana* (Cook 2019), historical accounts of other flying Strepsiptera, and photographic evidence, I conclude that the best way to characterize strepsipteran flight is through wing and power loading.

### Wing loading

Wing loading is total weight divided by wing area.^6^ It is important in quantifying maneuverability and speed of flight. For a given weight, the larger the wing area, the lower the wing loading. Low wing loading is associated with high lift, high maneuverability, and low power requirements, but also reduced speed (Cavagnaro 2019). High wing loading supports high sustained flight speeds and less perturbation by wind gusts, but requires more power, provides less maneuverability, and increases the difficulty of becoming airborne (Cavagnaro 2019). Tradeoffs are made between these traits based on the phylogenetic heritage and ecological needs of different flying animals.

### Flight muscle ratio

The flight muscle ratio (FMR) is the proportion of body mass a volant animal dedicates to flight muscle (Marden 1987). The higher the FMR, the more vigorously an animal can fly, but at the cost of whatever else the flight muscle mass might have been used for.

### Clap-and-fling

Clap-and-fling is a lift-enhancing mechanism used especially by small insects (Weis-Fogh 1973; Wootton 1990), including Strepsiptera. In it, the wings are struck together, generally at the top of a wingstroke, canceling the oppositely circulating air currents attendant to the wings. This frees them to immediately adopt new circulations (Chin & Lentink 2016), rather than having to wait for the preexisting ones to separate from the wings. The wings are then rapidly flung or peeled apart, generating strong oppositely circulating vortices (thereby observing the conservation of angular momentum) at the leading edges of the wings (Lehmann 2004). Thus, lift is generated more quickly and over an extended range (Chin & Lentink 2016), producing additional unsteady aerodynamic effects that result in nearly ¼ more lift than conventionally flapped wings (Marden 1987) (which are also “unsteady” in that they are flapped, rather than remaining essentially motionless, as in fixed-wing aircraft).

Marden (1987) also found that maximum liftable weight loads scale as the 1.0 power of muscle mass. Therefore, no matter how much muscle a flying animal has, having a given amount more would allow it to do the same amount more work (although the cost of doing so could also increase; e.g., the exoskeleton may not be able to support the addition muscle). Because all flight muscles output similar power per unit mass, FMR is an excellent stand-in for power loading when working with animals. However, power loading applies to both animals and constructed flying objects.

### Power loading

Power loading is weight divided by power—the rate at which energy is converted to useful work. With increased power (i.e., *lower* power loading) insects can fly faster, so low power loading is often desirable. However, increased power also increases energy demand. Because asynchronous flight muscles work most efficiently at a more or less constant resonant frequency (Josephson et al. 2000) when the wingstroke amplitude remains constant, most insects usually produce roughly the same power output regardless of actual demand. To allow slower flight, excess power is wasted, generally by lowering the body axis from horizontal (David 1978). In adult male Strepsiptera, there is little of the apparatus normally allocated to eating: the ventral mouthparts and their muscles are strongly reduced, the salivary glands, their receptacle, and the tentorium are all absent, and the midgut is strongly modified to hold air rather than food (Pohl & Beutel 2008), giving it the designation, “balloon gut” (Beutel & Pohl 2006). Accessory genital glands are also absent, and the Malpighian tubules are vestigial or strongly reduced (Pohl & Beutel 2008), all of which make for an expressly light frame that lowers power loading.

##### The mysterious balloon gut

The strepsipteran “balloon gut” almost certainly helps expand the bodies of teneral pharate adults by forcing hemolymph into compressed areas, as is common in insects through fluid ingestion (Jousset-de-Bellesme 1877; Prell 1914; Fraenkel 1935). This reserves space for enlarged flight muscles, etc., before the adult cuticle hardens. Anterior balloon gut extensions (Beutel & Pohl 2006) may provide thermal insulation, as thoracic air sacs do in dragonflies (Heinrich 1995). Finally, sensitivity to barometric pressure has been found in various bark beetles; a capacity linked to a large bubble in the otherwise evacuated ventriculus obtained by swallowing air prior to dispersive flight (Bennett & Borden 1971). It is hypothesized that atmospheric pressure-induced volumetric variation of the bubble allows the beetles to avoid flying on days with unstable weather (Bennett & Borden 1971; Lanier & Burns 1978). The balloon gut could confer this ability to adult male Strepsiptera.^7^

One function the balloon gut certainly does <u>not</u> perform is reducing flight weight. In an atmosphere, unpressurized air within a body weighs the same as the external air, and thus has no effect on overall weight. Actively ingested air, such as that sucked into the balloon gut by the strepsipteran air uptake apparatus (Beutel & Pohl 2006; Pohl & Beutel 2008), would be pressurized and would thus (v-e-r-y slightly) *increase* the insect’s weight. Even if the balloon gut were to somehow produce a vacuum, its limited volume would not result in much weight reduction at all (Gunn 1931). Appeals to relative density (specific gravity) fare no better: despite a reduction in overall density from increasing volume by ingesting air, the insect would still weigh the same (probably somewhat more, due to pressure), so the same force would be required to lift it (Gunn 1931). Thus, the balloon gut itself cannot be said to lighten Strepsiptera, although it does allow them to have the same relative volume as if they had a functional digestive tract, without the weight associated with it.

### Strepsipteran flight muscle ratio

I am unaware of the FMR ever having been determined for any strepsipteran species. However, for these considerations, it is not necessary to know the actual FMR; only that it is substantial. Fortunately, there are several indications that that is the case.

First, all known Strepsiptera are utterly dependent on optomotor anemotaxis (flight in wind to track sex-pheromones) for propagation. Second, species infecting stylopized Andrenidae are able to fly early in February in Germany, when even their much larger host bees are still lethargic (Ulrich 1933). The ability to fly over a wide range of temperatures, and especially at low thoracic temperatures, is associated with strong and versatile fliers (Krogh & Zeuthen 1941; Marden 1994). Third, in early spring in northern California, *Stylops pacifica* typically fly in sustained winds of 5–10 mph (8–16 kph), sometimes with gusts estimated as high as 25 mph (40 kph) (Linsley & MacSwain 1957). That a 2.75 mm (Bohart 1936) insect is able to fly in such conditions is further evidence of a well-developed flight apparatus. Finally, there are accounts of high speeds at which Strepsiptera have been noted to fly (Hubbard 1892; Pierce 1909)—a direct indication of high FMR.

### Strepsipteran flight

Strepsiptera have low wing loading (highly maneuverable, generally slow flight), but also high FMR (ample power compared to body weight, enabling great force production and acceleration). The application of these attributes to sex pheromone tracking accounts for their wide variance in flight characteristics.

## Methods

### Equipment used

Most photographs were taken with a Canon EOS RP (Canon Inc.; Ota City, Tokyo, Japan). I previously used a Canon T3i (aka EOS 600D) and briefly rented a Canon 5DS. After obtaining a Laowa 100mm F2.8 CA-Dreamer Macro 2X lens (Venus Optics; Hefei, Anhui, China), I used that lens exclusively. Each evening, a camera was mounted on a horizontally leveled, self-modified Cognisys High Speed Insect Capture System (now discontinued). All images were collected in August of 2019–2021 from several backyards in in Norman and Noble, Oklahoma, on nights with nearly still air, as is typical for the area. Images were mainly processed using Affinity Photo 1.10, and later 2.5.3 for wing area calculations.

### Obtaining photographs of flying Triozocera texana (née T. mexicana)

Macrophotography of flying insects remains largely confined to indoors, although current technology allows it to be extended to the field, where it is well-suited to recalcitrant insects. However, a camera’s built-in shutter is far too slow to capture clear images of quickly flying or tiny insects (magnification increases effective velocity). Since *Triozocera* are nocturnal, I avoided the nuisance of an external highspeed shutter because darkness itself can act as one (McIlleron & de Moor 2011). Thus, the camera was repeatedly placed into Bulb mode for up to a 2 minutes before noise accumulation became limiting.

Shepard (1979) identified the start and duration of the mating season of *Triozocera mexicana* (Strepsiptera: Corioxenidae), now *T. texana* (Cook 2019). I verified their continued local presence by investigating Norman Mosquito Authority (NMA) by-catch. The NMA subsequently escorted me to several of their most Strepsiptera-productive light traps, from which I inferred preferable host habitat. I used a New Jersey light trap (minus killing agent) to determine that male *T. texana* fly from about midnight to shy of 7 AM on weather permissive nights. I checked the first and a subsequently added (and differently located) trap hourly, whereupon several males could still fly. This occurred repeatedly, indicating trap placement in areas regularly traversed by mate-seeking males. To photograph Strepsiptera, I replaced one of the light traps with my insect rig. Because I knew adult males would be strongly drawn to UV (James & Strong 2018), on the rig I set a UV light above the focal point of a lens carefully focused on the intersection of a pair of laser beams. The beams were red and thus invisible to Strepsiptera. However, red is very visible to cameras, so the laser intensities were set extremely low, which also enabled tiny insects to easily trip the system. Whenever triggered by a disoriented insect flying through one or both laser beams, a pair of rig-mounted flashes fired, the shutter was closed, and an image was recorded.

Some 50 different flying males were photographed in this manner. At no point did I have access to a flight chamber, particularly not in the field, and certainly not during the COVID epidemic, when most of this work was done. Therefore, it was not possible to also transfer active males from a killing-agent-free light trap, to a flight chamber.

### Wing loading data sets

I combined two data sets containing measured insect wing loadings: Byrne et al. (1988) and Tercel et al. (2018). Byrne et al. (1988) were particularly concerned with Aphididae (aphids) and Aleyrodidae (whiteflies), both tiny insects in order Hemiptera, and thus not well-represented in previous data sets. They therefore amalgamated their own data with information collected from earlier studies. Tercel et al. (2018) were primarily interested in wingbeat frequencies, which in former meta-analyses had been collected at different temperatures using multiple methodologies. They compiled their own data set under standardized conditions, but subsequently discovered that order-level phylogenetic patterns were consistent with preexisting studies (Tercel et al. 2018). Consequently, I combined their data set with the data set of Byrne et al. (1988) to provide more complete coverage. Byrne et al. (1988) had already averaged multiple readings from singular species whenever they had been collected by the same research team. I expanded that to cover all entries, except when sexual dimorphism explicitly factored in or appeared to have done so.^8^ Therefore, apart from known sexual dimorphism and two other exceptions, each species is represented only once in the final compilation. Together, the combined data set contains 11 insect orders and 234 entries (231 species). However, four of the orders—(1) Blattodea, (2) Ephemeroptera, (3) Trichoptera, and (4) Mecoptera—are only represented by a single species each, and the first three, by a single specimen. Unfortunately, Strepsiptera were not included at all.

### Estimating strepsipteran wing loading

Calculation of an insect’s wing loading requires the fresh (wet) weight of its body and its wing area.

### Calculating wing areas

I detached the wings of several *T. texana* specimens preserved in 70% ethanol, spread them out flat, and photographed them alongside a paper ruler to calibrate the camera’s pixels when calculating areas. However, the wings were so delicate I only succeeded in doing so without complication twice, and unfortunately in one such case I had previously beheaded the specimen to prepare its eyes for electron microscopy. In four other instances, I digitally altered or combined torn wings to arrive at complete wings, but only used those for comparison. Instead of summing the areas of the two wings of a single specimen, the area of a single wing was doubled. All sampled wings and bodies appear in Fig. S1.

### Estimating wet weight

Sage (1982) produced several models to predict the wet and dry weights of various groups of insects. I used the Diptera-Hymenoptera model (R^2^ = 0.94; certified for body lengths 2.5–21.8 mm (Sage 1982)) to estimate *Triozocera texana* wet weight because it includes nematoceran flies, which Strepsiptera resemble. However, the only *T. texana* for which I had the wing area and intact body was 2.38 mm long, 0.12 mm below the certified range. To accommodate the model, I used the weight estimate for a specimen 2.5 mm long. I also calculated a hypothetical strepsipteran wing loading based on the wing area of *T. texana* and the average mass of the much larger *Stylops ovinae* (ca. 4 mm) (Pohl & Beutel 2013; Pohl et al. 2020). Both should produce high estimates. But if the wing loadings are nonetheless low, it can be concluded that the *T. texana* wing loading is also low, and by extension, that of Strepsiptera.

## Results

Tarsi of extant Strepsiptera only end in well-developed claws in the basal families Bahiaxenidae and Mengenillidae (Bravo et al. 2009); in all others, they are either reduced (Corioxenidae), or absent altogether (Pohl & Beutel 2008). Thus, in the following photographs, it is common for the tarsi to be tipped with seeming yellow or white balls. These are adhesive (tenent) setae (Pohl & Beutel 2008) that better equip the legs to adhere to appropriate areas of the host (Pierce 1909; Pohl & Beutel 2005, 2008) and to each other when grasping, but perhaps to little else (Pierce 1909; Linsley & MacSwain 1957).

### Wing loading

As discussed in *Methods*, the body length of the test specimen was 0.12 mm below the lower range for which Sage (1982) verified his model, but to satisfy model constraints, I submitted a body length of 2.5 mm. This should result in an overestimation of the weight of a 2.38 mm *Triozocera texana*. If, despite this, the wing loading is still low, *T. texana* should be considered to have low wing loading. The wing area for the *T. texana* specimen was (0.03534 × 2 = 0.07068 cm^2^). Applying Sage (1982) to a 2.5 mm long specimen produced an estimated weight of 0.00103 g (1.03×10^−3^ g), and a wing loading of 0.0146 g/cm^2^. In the combined data set of increasing wing loading from Tercel et al. (2018) and Byrne et al. (1988), *T. texana* sits between 48 and 49 of 231 surveyed species, with wing loading higher than *Myzus persicae* (green peach aphid), but lower than *Aglais io* (peacock butterfly). Intriguingly, if the average of all six of the single wing areas from Fig. S1 is doubled and divided into their estimated average weight (Sage 1982)—including three specimens below the lower length cutoff of 2.5 mm (i.e., 2.22, 2.3, and 2.38 mm)—the resultant wing loading is also 0.0146 g/cm^2^.

Among Lepidoptera, an insect order renowned for its low wing loading, this ostensibly over-weighed *T. texana* slots in between entries 22 (0.014 g/cm^2^) and 23 (0.0147 g/cm^2^) of 68. Therefore, its wing loading is lower than more than ½ of all surveyed moths and butterflies—it sits right at the cusp of the lowest ⅓. The lowest surveyed lepidopteran wing loading was that of *Pieris napi* (0.004 g/cm^2^); the highest, the sphingid moth, *Madoryx oiclus* (0.360 g/cm^2^).

Even calculating wing loading based on the wing area of the 2.38 mm *T. texana* specimen and the measured average wet weight (1.96 mg) (Pohl et al. 2020) of much larger (≥ 4 mm) *S. ovinae* results in a relatively low wing loading of 0.0277 g/cm^2^. This sits between entries 78 (*Tipula* sp., an unspecified crane fly) and 79 (*Sympetrum meridionale*, the southern darter dragonfly) of the combined data set, producing a wing loading that is less than nearly ⅔ of all surveyed species, and less than that of more than half of the surveyed moths. Thus, Strepsiptera have low wing loading.

The 4 megadiverse insect orders were among the 5 orders with highest average wing loading, ranging from 0.177–0.0665 g/cm^2^. The standout, Blattodea (0.149 g/cm^2^), was represented by a single species. Surprisingly, Coleoptera have higher wing loading than Diptera (0.134 vs. 0.0677 g/cm^2^) (Byrne et al. 1988; Tercel et al. 2018), despite the slow (but bumbling) flight of beetles, and the fast (but nimble) flight of higher flies. Thus, wing loading alone is insufficient to adequately characterize insect flight.

### Power loading

“*Elenchus tenuicornis* is able to fly vertically from the puparial surface” (Varley and Kathirithamby) (unpublished), reported in Smith and Kathirithamby (1984). I witnessed a *Triozocera texana* I thought was exhausted regroup, take off vertically without jumping, and spiral up to my headlamp. Among the 49 insect species he studied, Marden (1987) found that vertical takeoffs only occurred in insects with FMRs of at least about 0.25, regardless of whether or not clap-and-fling was employed. Thus, Strepsiptera have a flight muscle ratio of at least 0.25. As discussed in the Introduction, Strepsiptera have several adaptations that reduce weight without reducing flight muscle mass, thus increasing the flight muscle ratio. Assuming these reductions (including, e.g., replacement of the alimentary canal with a tissue-free “balloon gut”) together amount to at least 10% of the total body mass, the FMR is increased to 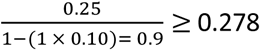. Furthermore, the strepsipteran flight muscle appropriation is known to be extensive (Smith & Kathirithamby 1984; Pohl & Beutel 2008; Fig. 2). If strepsipteran flight muscles are assumed to be at least 10% larger than normal,^9^ then the total FMR would be at least 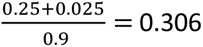. This should be considered a conservative estimate of the minimal strepsipteran FMR.

**Fig. 2.**
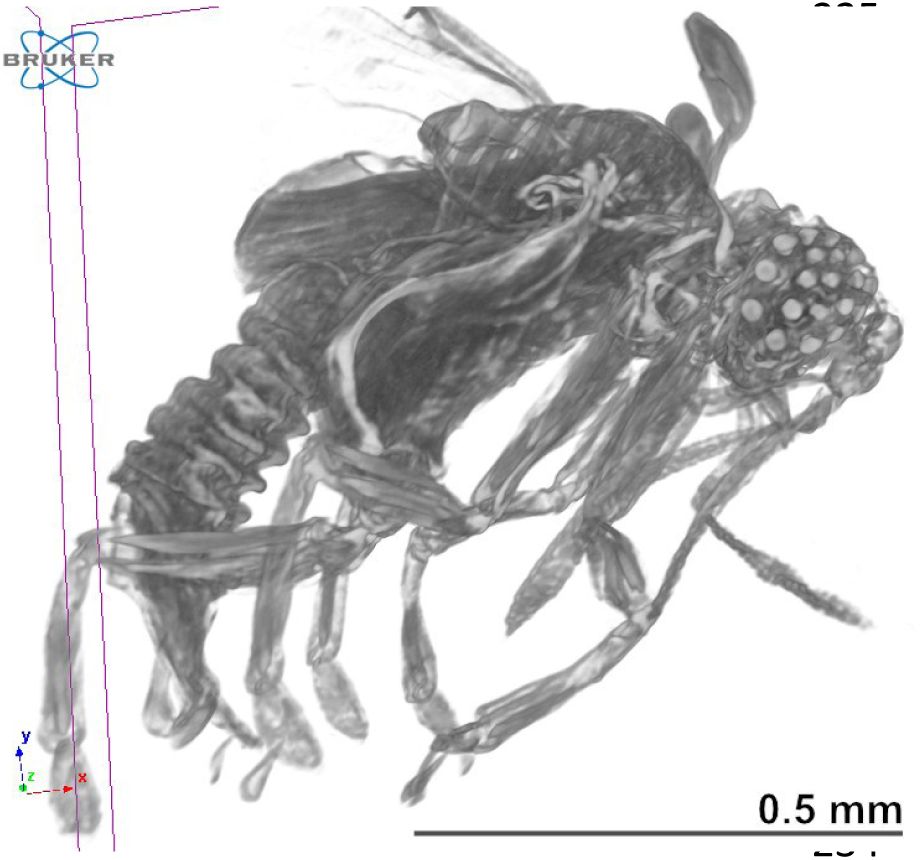
Flight musculature. Micro-computed tomography (micro-CT) image of an *Elenchus koebelei* showing the metathorax and a sketch of its internal flight musculature. The small hump behind the eyes is the mesothorax, from which the halteres emanate. The greatly expanded metathorax supports the only pair of wings. X-rays penetrate chitin, documenting that a large proportion of adult male strepsipteran anatomy is dedicated to primary flight muscles: Diagonal striations denote dorsoventral flight muscles (DVM) that extend from the oversized metaster-num (ventral sclerite of the metathorax) to the scutum, while horizontal striations indicate dorsal longitudinal flight muscles (DLM). Although the metathorax also contains the ancillary flight muscles that control wing orientation, they are overshadowed by the massive primary flight muscles. The protuberance extending from behind the notum (dorsum of the insect thorax) is the leading edge of the left wing. Scale bar approximate (body length of specimen was not recorded)

If 0.306 is accepted as the strepsipteran FMR, then according to Marden (1987), it would be nearly equal to the average FMR for Hymenoptera (i.e., 0.305). Thus, baseline strepsipteran FMR is higher than the average FMR for Hymenoptera. But beyond that, there is also the added efficiency of clap-and-fling.

That the proto-strepsipteran flight apparatus was already essentially that of extant lineages in all known strepsipteran stem groups (i.e., †*Protoxenos*,^10^ †*Cretostylops*, and †*Mengea*) is evidence that it developed early in strepsipteran evolution (Pohl & Beutel 2008). Moreover, a Burmese amber strepsipteran fossil, †*Cretostylops engeli*, provides strong evidence that the modern strepsipteran flight apparatus is at least ca. 100 myo (Grimaldi et al. 2005). Therefore, without contrary evidence, it should be assumed that all extant Strepsiptera have high FMRs, and thus low power loading.

### Wing displacement

One may conclude from Fig. 3 that among extant Strepsiptera (Pohl & Beutel 2008), whenever a wing’s leading edge is perpendicular to the body axis (Fig. 4a) or angled further back (Figs. 4c–g, 5e), there is either infolding (Fig. 6e)—typically at or near the wing-body interface (Fig. 4a&c)—or a portion of its leading edge is folded over (Fig. 6a). Extreme flexibility allows for this, which may itself result from the wings being teneral (Figs. 5f). Moreover, the shape of the metanotum in particular (Fig. 1), facilitates wing area reduction in flight, as does the upward curvature of the distal abdomen of Strepsiptera engaged in forward flight (Fig. 4c).

**Fig. 3.**
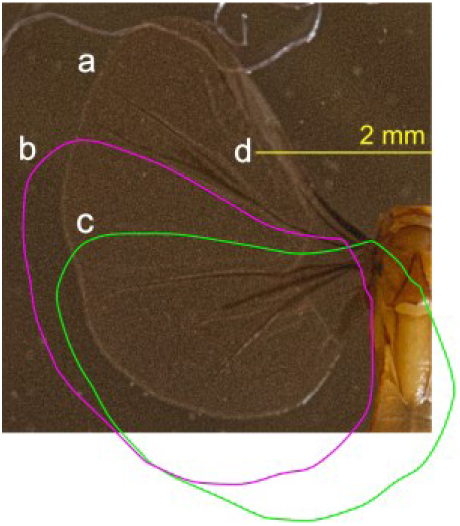
Headless *Triozocera texana* with attached intact left wing. (a) How far forward (60°) the outstretched wings extend without any wingtip folding or rotational displacement. (b) How far forward (30°) the outstretched wings extend when the lateral wing surface just touches the body. (c) How far into the body the wings protrude without any infolding or camber, when the wing’s leading edge is perpendicular to the body axis. (d) Farthest forward extent of antennaless head. (Head length approximated by fitting resized intact bodies to this headless body.)

**Fig. 4.**
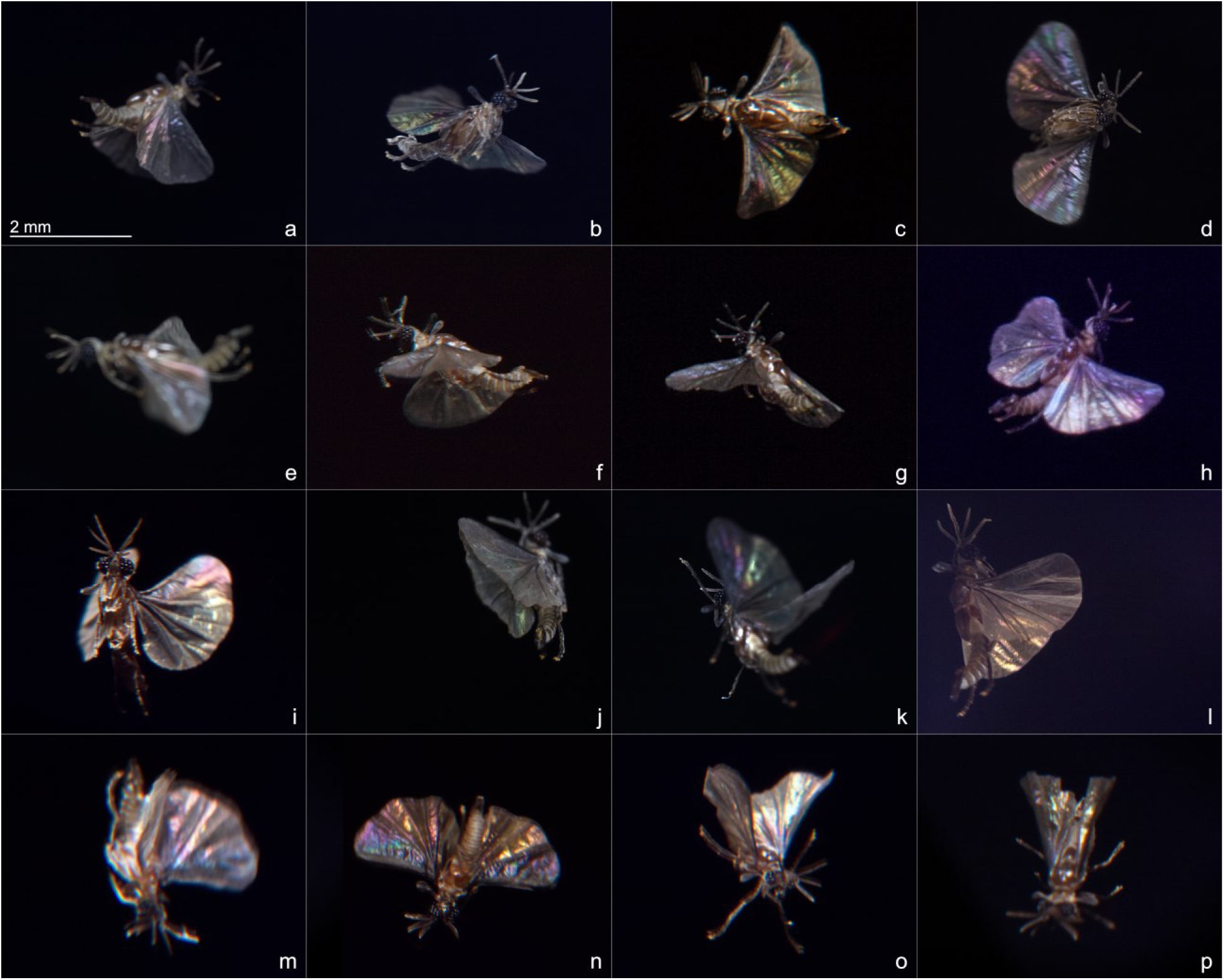
Potential strepsipteran flight modalities surmised from photos of them in flight in the field. *(a&b)* Cruising—body nearly horizontal with leading wing edges perpendicular to body axis or further forward: (a) upstroke (reduced wing area); (b) likely in downstroke (wing area expanded and not flaccid). *(c&d)* Rolled 90° (knife-edge maneuver): (c) cw; (d) ccw; changing trajectory (lower wing has greater tilt and blur than upper wing). *(e–h)* Reduced wing width: (e) wings displaced posteriorly; (f) leading edge of nearest wing visibly folded over; (g) same insect shot at same time as *(f)*, but at different angular offset; (h) forward edge of right wing visibly folded over. *(i–l)* Hovering—body nearly vertical, wings outstretched for maximum lift: (i); (j) clap-and-peel; (h) just after clap-and-peel on downstroke; (l) wings tilted inward at end of downstroke in “near” clap-and-fling, increasing lift. *(m&n)* Probable power dives: (m) wings engaged in clap-and-peel, legs held in close, head aligned with body; (n) different point in wingstroke than *(m)*. *(o&p)* Controlled descent—legs extended, head not aligned with body axis, wings passively modifying airflow: (o) wings fluttering; (p) wings pressed back. All insects presented at scale at 1.98× magnification, with possible exceptions of *(c*, *g*, and *h)*, whose original magnifications were estimated. *(a*, *b, d, e, j, k)* shot with Canon RP; *(f, i, m–o)* shot with Canon 5DS; *(c, g, h, l, p)* shot with Canon T3i. *(m–o)* were blurred due to poor flash synchronization

**Fig. 5.**
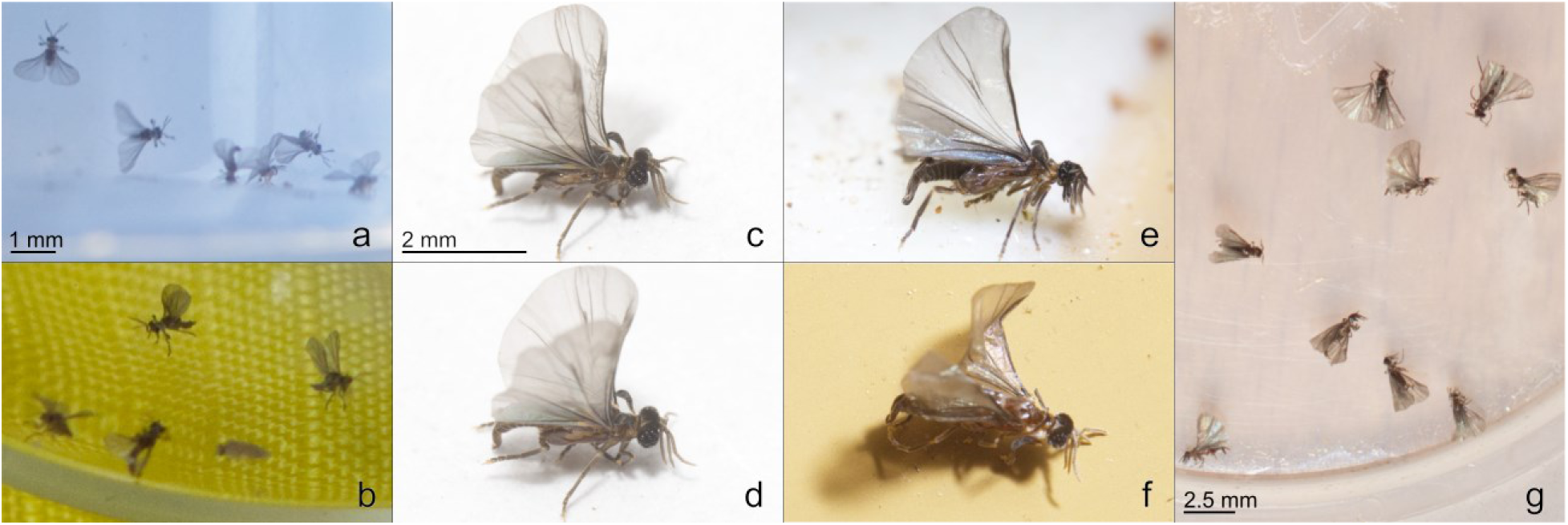
Resting wing positions. *(a*&*b)* are photos of *Elenchus koebelei* that could all still flap their wings, though most could no longer fly. *(c–g)* are photos of *Triozocera texana*. In *(c–e)*, *T. texana* were provided with areas free of other species. Alteration in wing position suggests they cannot lock their wings vertically. (a) *E. koebelei* wings are unpleated and comparably placed when oriented horizontally. (b) Two *E. koebelei* have their wings outstretched vertically; however, they are held apart unlike when they can no longer flap them. *(c&d)* are photos of the same *T. texana* taken at slightly different times. (c) The wings are held vertically, roughly perpendicular to the long axis of the body. (d) The wings are positioned vertically, but have been pulled forward and project laterally slightly. (e) The wings are held vertically, but have been pulled back somewhat. *(f&g)* Unlike the others, these insects were already dead. (f) This specimen expired on the insect rig itself. Its wings subsequently adopted the form of the edge on which it had died. This is a strong indication that upon emergence strepsipteran wings have not actually hardened. (g) Images of several specimens collected one evening. Their wings are oriented haphazardly: one horizontally, some vertically, several intermediate between those extremes—as if upon death, the wings dried completely in whatever position they occupied when the insect depleted the strength to move them. Scale bars are reasonable estimates that remain in effect until alphabetically superseded

**Fig. 6.**
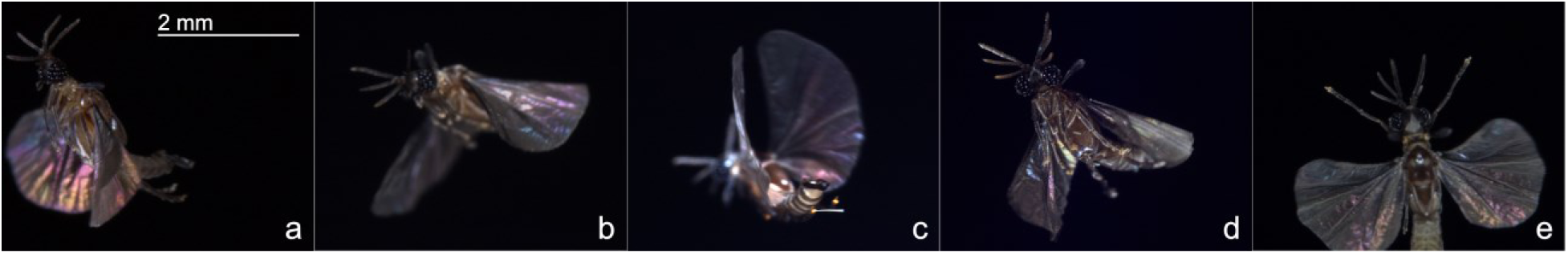
Hallmark strepsipteran wing pliability. (a) Despite the rigidity associated with leading edge spars, here the foremost wing area is flaccid. *(b&c)* Wings flex up and down, allowing both (b) positive camber, and (c) negative camber. (d) Wings folded up like a fan, perhaps in transition. (e) Strepsiptera may partially collapse their wings on the upstroke

### Strepsipteran flight versatility

Figure 4 is an overview of photographically captured potential strepsipteran flight modalities, including level cruising (Fig. 4a&b), the knife edge maneuver (Fig. 4c&d), reduced wing width (Fig. 4e–h), hovering (Fig. 4i–l) (the closer to vertical, the slower the flight speed), clap-and-fling variants (Fig. 4j–m), powered diving (Fig. 4m&n) (Also see Online Resource 1), and controlled descent (Fig. 4o&p). In controlled descent, the specimens appeared to be looking at the camera lens (note the head orientations). However, Strepsiptera may actively dive to avoid echolocating bats.

Concerning controlled descent, in Fig. 4o, the wings were open wide enough to be flapped, but appear flaccid and tilted, rather than swept, back, and may have been disengaged to slow descent. Additionally, the legs are outstretched, further increasing drag. In Fig. 4p, the wings are drawn in too close for flapping, so descent was likely also passive.

### Reducing exposed wing area

Strepsipteran wings are very broad, limiting flight speed and increasing energy consumption. However, in Fig. 4f–h, the wings are folded over, probably in the upstroke, reducing wing area and thus drag. In Fig. 4e, the wings are either folded under or have been actively swept back—i.e., the wingtips may have been temporarily displaced posteriorly—which would increase wing loading in the power cycle of the wingstroke. This is known to occur with increased flight speed in the hawk moth, *Manduca sexta* (Willmott & Ellington 1997). Thus, folding or sweeping the wings back represents a potential mechanism by which Strepsiptera could produce and sustain higher flight speeds.

### Wing pliability

Strepsipteran wings exhibit remarkable flexibility (Fig. 6a–e), including the ability to fold over along the leading edge (Figs. 6a; 4f–h), positive and negative camber (Figs. 6b&c; 4d), and even complete collapsibility (Fig. 6d), in which the wings provide no lift to the airborne insect. Wing collapse may form a transition between flight modalities during unexpected events. To reduce air resistance, exposed wing area can be lessened during the upstroke (Figs. 6e; 4a).

## Discussion

**Table 1.**
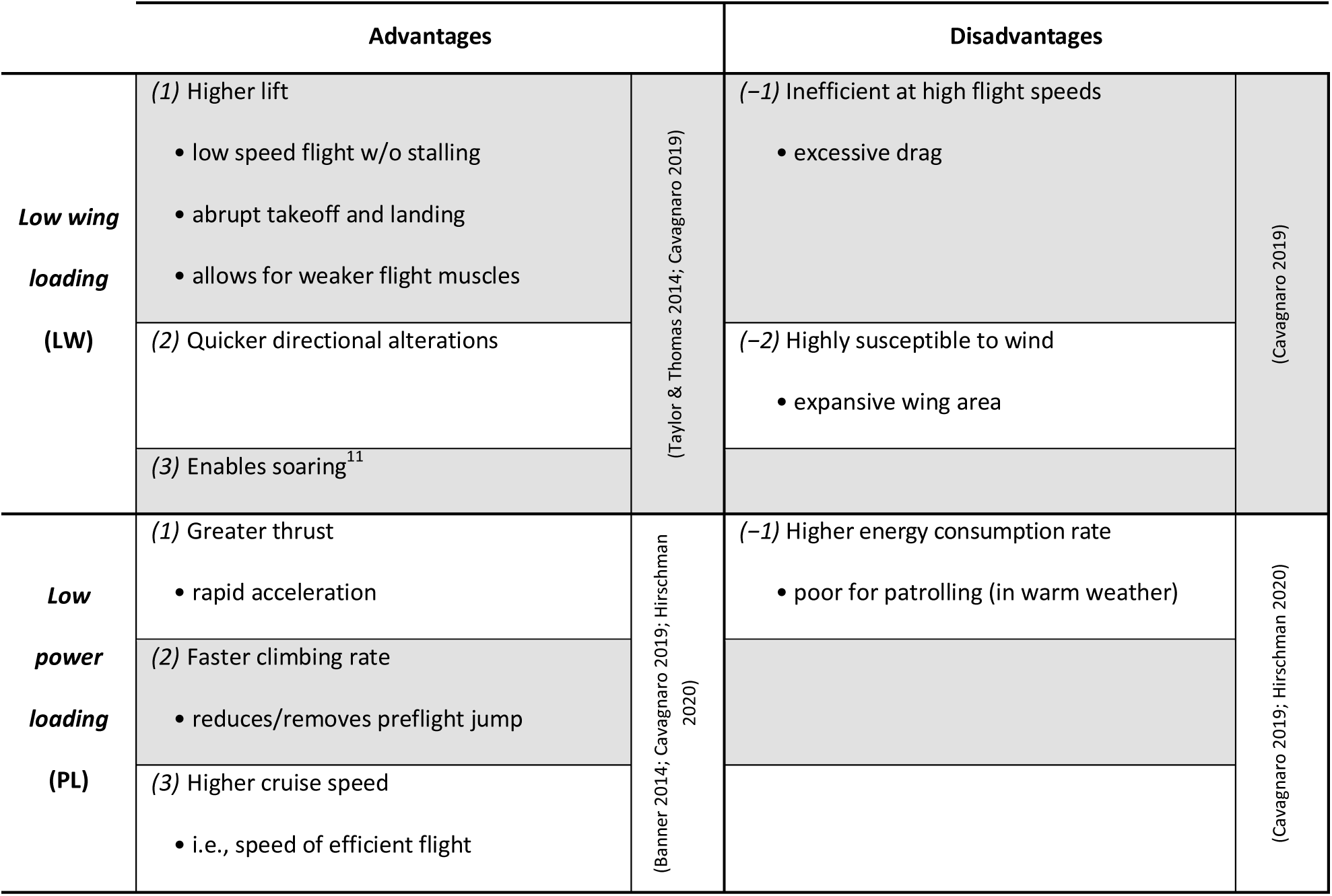
Low wing loading and high FMR together enable heightened aerial performance.

|  | Advantages |  | Disadvantages |  |
| --- | --- | --- | --- | --- |
| **Low wing loading (LW)** | (1) Higher lift<br>• low speed flight w/o stalling<br>• abrupt takeoff and landing<br>• allows for weaker flight muscles | (Taylor & Thomas 2014; Cavagnaro 2019) | (-1) Inefficient at high flight speeds<br>• excessive drag | (Cavagnaro 2019) |
|  | (2) Quicker directional alterations |  | (-2) Highly susceptible to wind<br>• expansive wing area |  |
|  | (3) Enables soaring<sup>11</sup> |  |  |  |
| **Low power loading (PL)** | (1) Greater thrust<br>• rapid acceleration | (Banner 2014; Cavagnaro 2019; Hirschman 2020) | (-1) Higher energy consumption rate<br>• poor for patrolling (in warm weather) | (Cavagnaro 2019; Hirschman 2020) |
|  | (2) Faster climbing rate<br>• reduces/removes preflight jump |  |  |  |
|  | (3) Higher cruise speed<br>• i.e., speed of efficient flight |  |  |  |

### Low wing and low power loading combined

The higher the wing loading (i.e., the smaller the wings or greater the weight), the harder an animal must work to remain aloft. But it is also generally true that the higher the wing loading, the faster an airborne animal is able to fly, especially for extended periods. Low wing loading optimizes for low-speed, highly maneuverable flight. High wing loading optimizes for high-speed, predominately straight flight.

Although beetles have even higher wing loading than flies, to account for their poor flight performance, Coleoptera must also have low FMRs (i.e. *high* power loading—high body mass compared to flight muscles mass). It is likely elytra provide lift enhancement (Le et al. 2014), allowing beetles to fly with even less power. Several beetles also exhibit clap-and-fling, some on both ends of the wing cycle (Yang et al. 2024; Fig. S3). In contrast, many Diptera must have high FMRs in addition to high wing loading to support their extremely nimble and rapid flight. Outstanding aerobatics despite high wing loading suggests that power output can be quickly and independently regulated between the two wings. This ability coupled with fast precise feedback from the halteres is likely what gives many flies their great aerial maneuverability. Strepsiptera have much lower overall wing loading than Diptera (< 0.0146 vs 0.0677 g/cm^2^), but it is reasonable to expect they are capable of similar feats when flying at high speed (e.g., (Hubbard 1892); (Pierce 1909, p. 14)), when wing loading can ostensibly be increased by sweeping the wings back (Fig. 4e) or allowing the leading edges to fold over (Fig. 4f–h).

### Elucidating diametric flight descriptions

#### Carried along by a gentle breeze

The “exceedingly graceful” flight observed by Thwaites (1841) is largely an expression of high lift and perhaps also high maneuverability, depending on the character of the gracefulness. Both are aspects of low wing loading (WL 1&2). Such flight is especially associated with males patrolling nest sites for stylopized *Andrena* in temperate climates in late winter and early spring (Friese 1883; Ulrich 1933).

#### Flight so rapid the eye can hardly follow

Based on the comparatively large volume of air strepsipteran wings displace with each wingbeat (Fig. 4), along with a respectable wingbeat frequency (150–160 Hz in tethered *X. vesparum* (Pix et al. 1993)), Strepsiptera are capable of quick acceleration. The “whirligig” flight Hubbard (1892) observed stemmed in part from this (see PL 3), and the ability to abruptly change direction (WL 2). It is plausible Hubbard’s *Xenos* sp. also increased their wing loading by sweeping their wings back (Fig. 4e) or folding the leading edges over (Fig. 4f–h). But even if so, rapid flight exacts an energetic toll (see WL −1).

#### Not so weak and helpless

A *Hylecthrus rubi* used its wings to drag around its dead host and two unemerged male conspecifics (SS Saunders 1853). That display of strength was related to low power loading (PL 1). Normal strepsipteran wingstroke amplitude is likely about 160° (Pix et al. 1993; Fig. S4), potentially reserving some 20° for responding to wing injury (Muijres et al. 2017), or maximizing force or speed.

#### Very peculiar unsteady flight

Sometimes Strepsiptera fly erratically in open air (E Saunders 1888), when it is energetically costly (WL −1) and without apparent payoff. In moths, losing a strong scent plume (normally indicating proximity) before establishing visual contact with a potential odor source can result in much higher flight velocity (PL 1) and wide fluctuations in direction (Baker et al. 1976; Mafra-Neto & Cardé 1994) (WL 2). Thus, casting as found in moths, seems to be what E Saunders (1888) observed and what Enock (1875) described as flight “not keeping in one direction more than a few inches,” albeit magnified by haltere-assisted aerial agility and expressly high FMR.

### Strepsipteran pheromone pursuit

There are two known strategies volant insects use to follow odor plumes (Dindonis & Miller 1980; Bursell 1984; Cardé & Willis 2008): Aim-then-shoot and optomotor anemotaxis. In aim-then-shoot, the current odor-bearing wind direction is assumed to be that of the odor emitter. That direction is determined before take-off and maintained while aloft, and backtracking or reassessment is only made if the odor is lost before its source is identified (Bursell 1984). If a potential odor source is not visually identified in a flying bout, an upwind-flying insect typically overshoots the odor emitter, lands, turns 180°, reassesses the wind direction, and flies downwind, but not as far, closing in on the target by repeating the process a few to several times. Because insects employing aim-then-shoot do not seem to regulate their ground speed when surging forward (Bursell 1984), they probably use skylight polarization to ensure straight flight, regardless of altitude and fluctuating wind direction. However, there is no indication that Strepsiptera have a dorsal rim area; moreover, none of their photoreceptor axons are known to proceed directly through the lamina to the medulla for polarization assessment. Aim-then-shoot is best known from a few Diptera (Dindonis & Miller 1980; Bursell 1984; Cardé & Willis 2008) (Cardé & Willis 2008), none of which follow odorants emitted from individual insects. Finally, aim-then-shoot requires landing and reorientation on potentially unfamiliar substrates, for which Strepsiptera are ill-equipped, due to their specialized tarsi (Linsley & MacSwain 1957; Pohl & Beutel 2008) and poor walking ability (Hubbard 1892; Linsley & MacSwain 1957).

In optomotor anemotaxis, an insect—e.g., a moth—follows odor plumes upwind (anemotaxis) using the speed and direction at which visual stimuli cross its eyes (optic flow) to evaluate progress and to correct for deviations in intended flight heading (Cardé & Willis 2008). There are several flight patterns associated with pheromone-mediated optomotor anemotaxis (Traynier 1968; Murlis & Bettany 1977). These can be used to determine if salient features of strepsipteran flight are congruous with them.

**Table 2.**
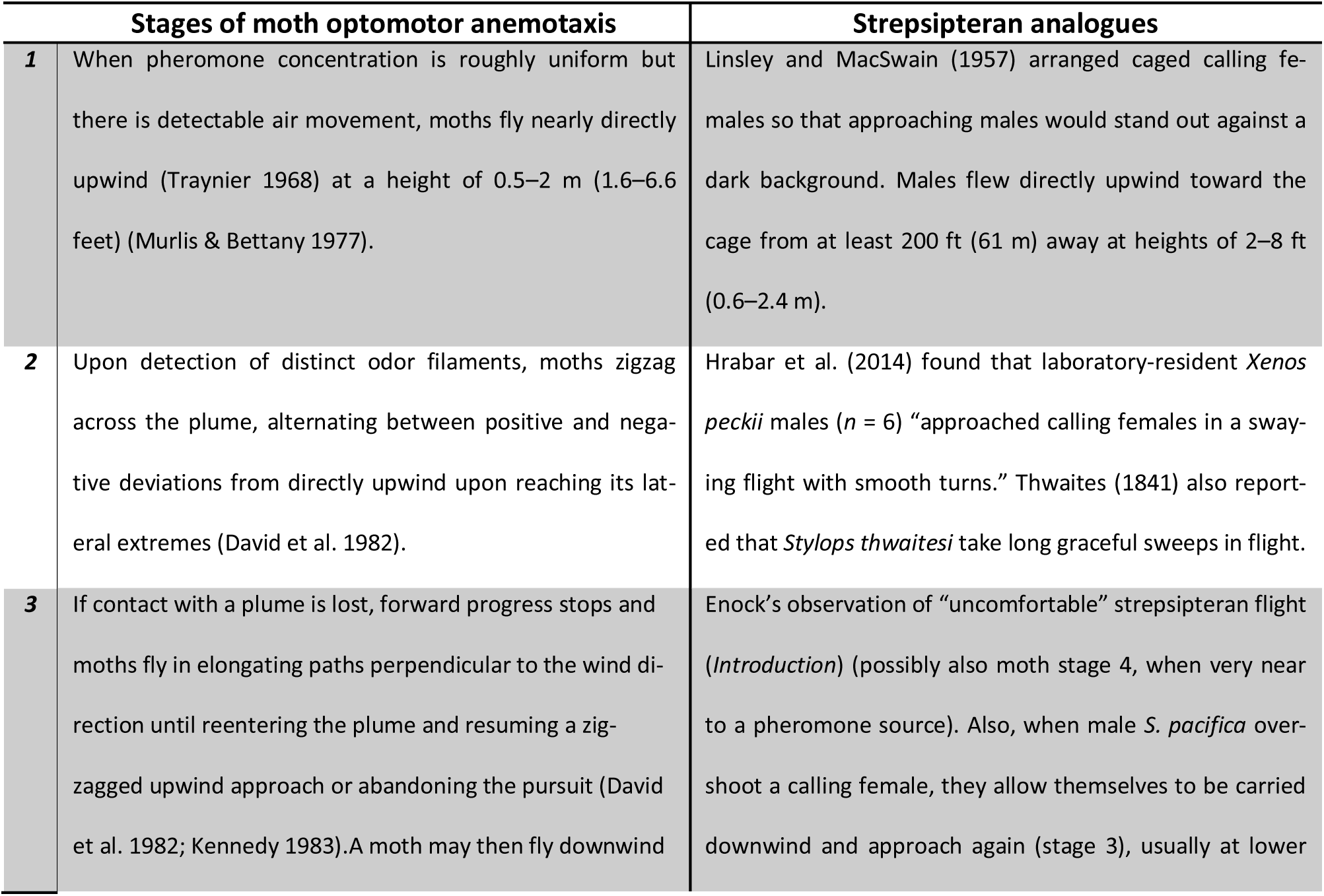
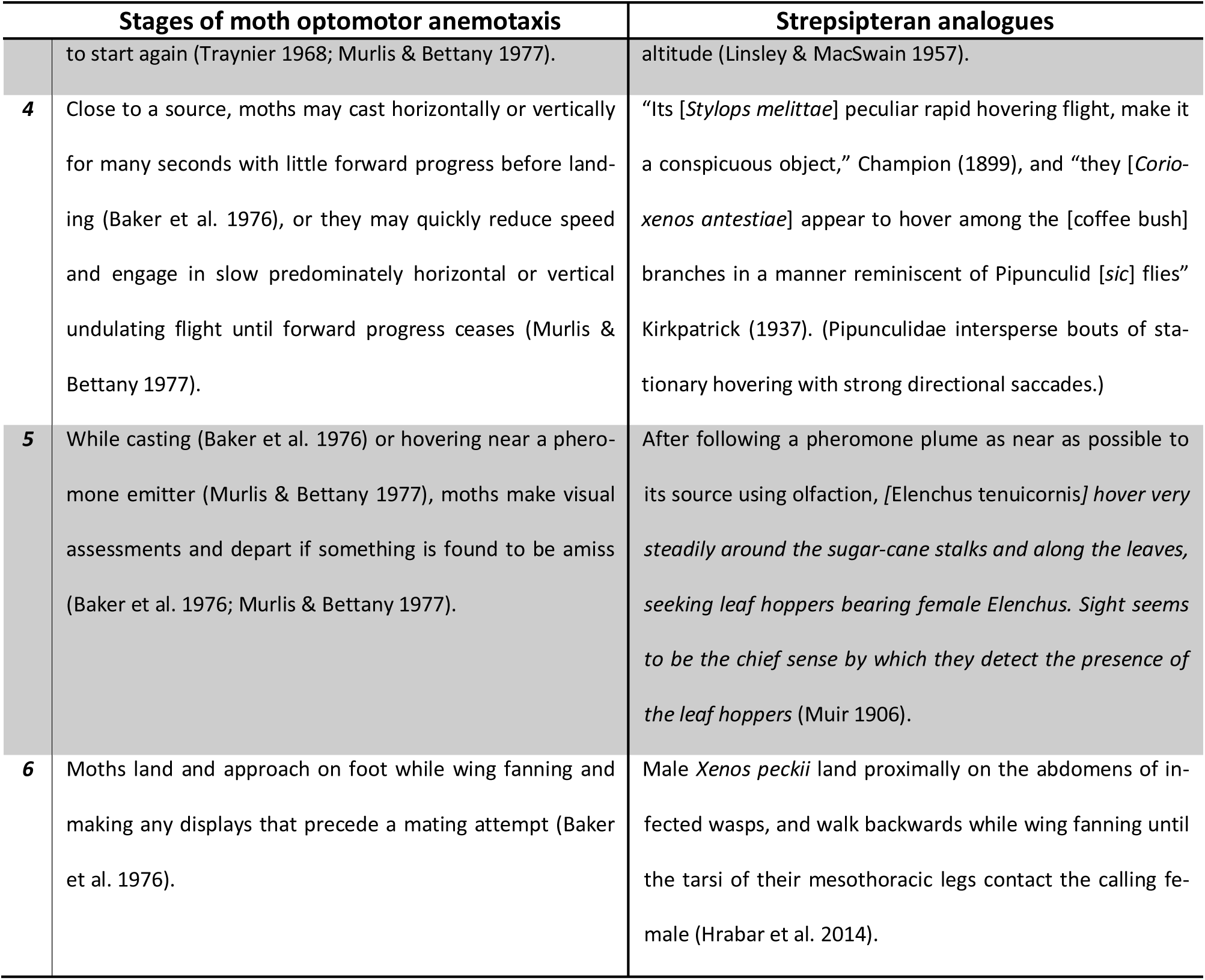
Recognized stages in moth optomotor anemotaxis compared with potential strepsipteran counterparts.

Note that wing fanning by male moths has been associated with sniffing, i.e., increasing airflow adjacent to olfactory surfaces to enhance smell (Baker & Cardé 1979; Loudon & Koehl 2000). Also, because Strepsiptera can hover (Fig. 4i–h), they can also fly backwards. From this and *Xenos* walking backward when approaching calling females, it follows that during such approaches, males reverse the airflow across their antennae with respect to its direction in normal flight.

### Explanation of strepsipteran flight

Reported instances of incongruous strepsipteran flight correspond directly to the major paradigms of sex pheromone pursuit found in moths and other insects. Furthermore, female Strepsiptera are known to produce sex pheromones (Cvačka et al. 2012; Tolasch et al. 2012; Zhai et al. 2016). Therefore, it is appropriate to conclude that adult male Strepsiptera use optomotor anemotaxis to trace sex pheromone plumes back to their sources.

Lepidoptera (Greenfield 1981), Neuroptera (Aldrich & Zhang 2016), and Trichoptera (Löfstedt et al. 2008) are also known use single-source sex pheromones. Each of these has low wing loading, but species from those orders may not have high FMRs. However, unlike them, Strepsiptera are parasitic, and furthermore, do not control the activity period of their hosts. The weight savings and extensive musculature Strepsiptera dedicate to flight are useful because eclosed adult males are tiny, are very likely to compete intensively for mates, have very broad wings whose effective area and length can be temporarily reduced, and have extremely short lifespan. Low wing loading allows adult male Strepsiptera to fly slowly (unlike a locust, e.g.) with high maneuverability (unlike most beetles), which is always beneficial for tracking single-source pheromone emitters. But high FMR also allows Strepsiptera to accelerate quickly and attain high relative flight speeds, improving the chance to exploit whatever access window the host exposes. High FMR also enables Strepsiptera to fly in cold weather.

Although Strepsiptera are capable of quick flight, they are usually encountered when flying slowly, first because it is easier to see them then; second, they fly closer to the ground when flying slowly, which also makes them easier to see; third, they tend to slow down around plants, where both infected hosts and potential observers are more likely to be. Strepsiptera typically sustain greater flight speeds higher above ground and farther away from calling females (Lindsay & MacSwain 1957), where odor plumes are less chaotic (Traynier 1968). Lastly, several *Stylops* sp. emerge in late winter or early spring (Ulrich 1933; Linsley & MacSwain 1957). Unsurprisingly, Strepsiptera fly slowly in such cold. Moreover, being one of the only insects active at the time, they are also more conspicuous.

### Why strepsipteran wings are as they are

As a rule, insect wings must be sclerotized before they are flightworthy (Prell 1914; Fraenkel 1935), but strepsipteran wings are an apparent exception (Fig. 5f). “Modern” Strepsiptera prepare their wings for flight while within pupal cases confined to the abdomens of other insects. This consists of wing inflation and elongation, but not spreading or sclerotizing, so wing folding cannot occur. Accordingly, strepsipteran wings have pleat affinities (Figs. 1 & 5), but no crossveins (Pohl & Beutel 2005) or fold lines. Fine, closely-packed, wing microtrichia (Fig. 7) probably reduce adhesion by resisting wetting (Polet et al. 2015) in humid puparia. The apparent thinness of strepsipteran wings increases flexibility, which may improve flight performance (Mountcastle & Daniel 2009; Fig 4f–h) apart from reducing wing weight. After adult emergence, the disadvantages of wings that cannot be folded are inconsequential for Strepsiptera: When not mating, or wing fanning in preparation for mating, eclosed adult males basically fly non-stop until they are unable to remain airborne.

**Fig. 7.**
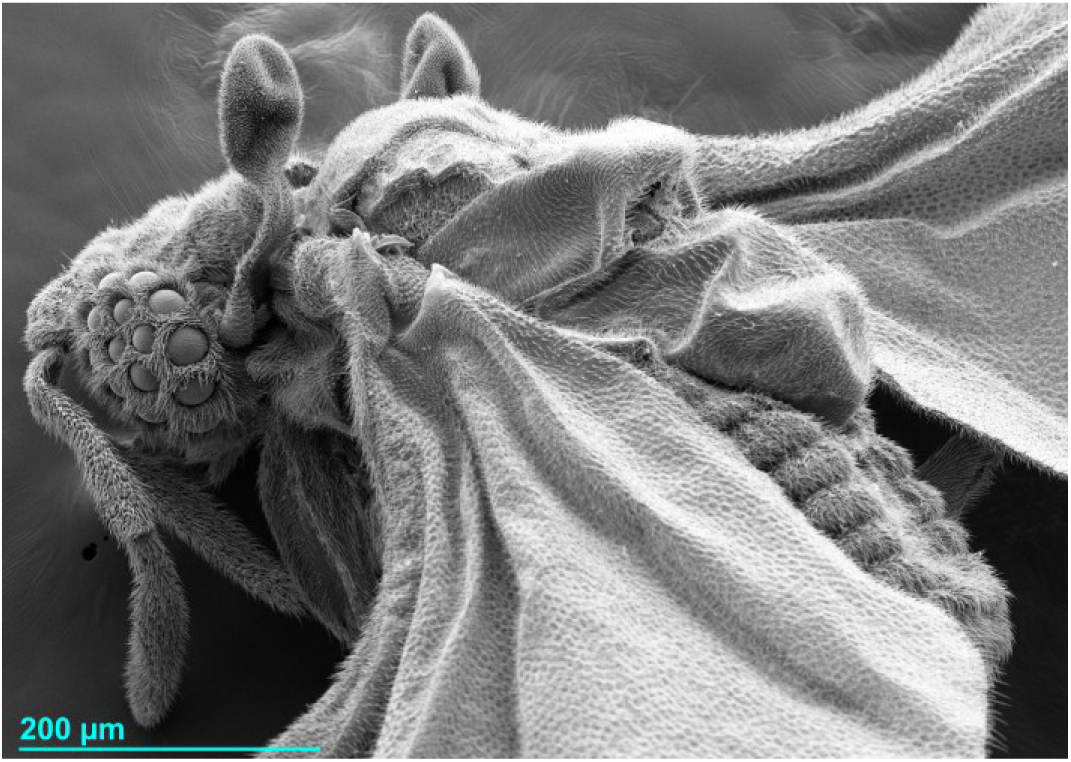
SEM image of an *Elenchus koebelei* at a magnification of 150×. Note the fine ‘hairs’ on the wings and most of the rest of the body. Strepsipteran microtrichia may be of particular importance for allowing the wings to be opened enough to expand longitudinally in the puparium during development—or else to at least be expanded while emerging. Also displayed is the enormity of the dorsal extent of the flight muscle-filled metathorax, as are the few large ommatidia.

### Flying on immature wings?

That the wings of an expired adult male adopted the shape of the object it died on (Fig. 5f) is a strong indication that the wings were teneral, even though they were used to fly to the site. Considering their extreme flexibility (Fig. 6a&d), strepsipteran wings may be teneral for all of an adult male’s active life (Fig. 5). Moreover, wings that are being cleared of the cells that secreted them are cloudy (Tögel et al. 2008). Intriguingly, strepsipteran wings have often been described as milky (Bohart 1936) or milky white (Curtis 1829; SS Saunders 1850; Pierce 1909; Smith & Hamm 1914), as if they had not been cleared at all, or as smoky hyaline, pale hyaline (Perkins 1905; Brues 1903), or subhyaline (Brues 1903), as could occur during clearing. Occasionally, the wings are simply described as hyaline (Pierce 1909), as might occur after clearing is complete. Thus, Strepsiptera likely fly on immature wings.

### Wing desiccation

Prior to emergence, the humid pupal chamber prevents desiccation in the absence of wing motion. But in both *E. koebelei* and *T. texana* that had been given free areas in which to walk, but had starved past the point of incessant wing flapping, the wings gradually set; most often in a vertically extended position (see Fig. 5). Thus, the wings either dry rapidly when hemolymph is not continuously pumped into them (Wootton 1992), or they gradually desiccate or sclerotize as eclosed adult males age. Flapping (Salcedo & Socha 2020) or wing fanning may help pump hemolymph, which would explain why Strepsiptera keep their wings in nearly constant motion for their entire active adult lives.

### To have or lack halteres

Low wing loading with modest power already provides high maneuverability. So why do Strepsiptera have halteres? Ostensibly, they had no primordial need to avoid predation. Fossils and extant basal species strongly suggest Strepsiptera were always small and originally nocturnal (Pohl & Beutel 2008). There are still no nocturnal aerial visual predators, insect or otherwise, and according to existing fossil evidence, Strepsiptera evolved halteres long before bats first appeared (Simmons et al. 2008; Speakman 2008; Misof et al. 2014). Modern Strepsiptera have few natural enemies (Muir 1906; Kirkpatrick 1937), and well-armed or potentially aggressive hosts—mantids, ants, wasps, and bees—were only first colonized relatively recently (Pohl & Beutel 2005). Only males attempting to mate elicit responses from the host species (Muir 1906; Smith and Hamm 1914; Hughes-Schrader 1924), and only attempting to mate at a nest site of eusocial Hymenoptera appears to provoke aggression (e.g., (Hubbard 1892) vs. (Brues 1903; Muir 1906; Hughes-Schrader 1924; Ulrich 1933; Kirkpatrick 1937; Linsley & MacSwain 1957)).

However, unlike low wing loading, halteres also provide feedback enabling insects equipped with them to directly correct orientation in response to perturbation. Because halteres do not rely on vision (Chan et al. 1998), they maintain functionality even if the visual system provides limited spatial or temporal resolution. In fact, based on their persistent presence in wingless (Dick & Pospischil 2014) and even blind bat flies (Theodor 1967; Reeves & Lloyd 2019), halteres are useful apart from flight, and also in the complete absence of eyes. Ocelli are faster than compound eyes, but halteres operate more quickly than either (Krapp 2009), making them especially valuable in low-light, when ocular input is delayed. The absence of ocelli and low acuity of strepsipteran compound eyes could help explain spasmodic flight in Strepsiptera, there being nothing to handle stimuli of intermediate speed. That and the combination of low wing loading and high FMR could make for exceptionally jarring flight, especially if juxtaposed against the patient graceful flight Strepsiptera are also capable of. Thus, halteres allow high maneuverability while flying rapidly, but ostensibly did not evolve for that purpose in Strepsiptera.

### Vision in Strepsiptera

Strepsiptera are well-known for having small numbers of very large ommatidia. Optomotor anemotaxis requires that a flying insect discern enough environmental features to determine its bearing (based on *how* the features move along its eyes) and the rate at which it is moving with respect to its surroundings (from *how fast* features move across its eyes), so an appraisal of strepsipteran vision is in order.

Ancestrally, Strepsiptera were tiny nocturnal insects (Pohl & Beutel 2008), so high resolution vision was implausible (Buschbeck et al. 2003). Despite an interesting foray (Buschbeck et al. 1999, 2003), there is no evidence of strepsipteran superommatidial resolution (so-called “chunk-vision”) in diurnal species either (Pix et al. 2000)—and also no need for it, since all of the problems it would address were already solved by nocturnal forebears. Instead, as non-visual sensors, halteres ostensibly mitigate low spatial resolution by enabling course correction without recourse to vision, while freeing space by making ocelli redundant. Moreover, mate-seeking Strepsiptera do not care *where* they are going, as long as it is likely toward a calling female, so the dorsal rim area has also been discarded, again enhancing vision by leaving more area for image perception. Adult males do not need to visually discern discrete environmental features; they only have to be able to tell them apart—something that great sensitivity seems to support in the absence of high resolution or acuity. Detected light levels ostensibly change when the position of a very sensitive eye is translated or rotated, which is apparently enough to provide reliable input for optic flow. It is likely for this reason that the eyes of diurnal Strepsiptera retain so many features normally associated with nocturnal vision, such as large ommatidia, reticulate retinas, and high sensitivity (Buschbeck et al. 1999; 2003).

#### No new visual paradigm

In basal Stylopidia—Strepsiptera in which adult females cannot depart the host—all of which were presumably nocturnal, exceptional mate recognition was achieved with minimal reliance on vision. Because these derived females are cryptic, there is no advantage to favoring vision in photopic environments. Thus, in diurnal species such as *Xenos peckii*—the very species for which Buschbeck et al. (1999) conceived of “chunk vision”—actual mate recognition is chemotactile. Males fly past calling females (from behind), land on the proximal end of the host’s abdomen, and walk *backward* along it until their middle legs touch the female, whereupon the protruding cephalothorax is immediately grasped, and mating begins (Hrabar et al. 2014). Moreover, another diurnal Strepsiptera, *Halictophagus silwoodensis*, was also found to chemotactilely identify females located on the underside of its leafhopper host, *Ulopa reticulata* (Henderickx 2008). Finally, Strepsiptera infecting bugs are hidden beneath their hosts’ hemelytra, and therefore cannot be visually detected at all (Kirkpatrick 1937). Thus, there is no need to evolve a massively more complex and expensive visual paradigm to partially and less robustly address a problem previously solved in its entirety.

Males of the extant pre-stylopidian families Bahiaxenidae and Mengenillidae are no larger than diurnal Xenidae, but have twice as many ommatidia. No one expects them to have “chunk vision,” but these nocturnal males do not have tarsal sensory patches, so their liberated female conspecifics *are* visually identified. Ironically, it was vision-based recognition that was superseded.

#### Can’t see the meadow (f)or the grass?

In taking flight from the ground in the midst of grass or other vegetation, males will frequently “fly-walk” up the stems, catapult into the air, collide with other stems, and fall to the ground a number of times before finally completing a successful take-off. (Linsley & MacSwain 1957)

Why would *Stylops pacifica* find it advantageous to catapult into the air when they already have low wing and power loading? Stiff breezes and wind gusts are frequent in early spring (Linsley & MacSwain 1957). Like *Lymantria dispar* (gypsy moths) (Murlis et al. 1982), *S. pacifica*, can likely orient and fly in higher winds than it can successfully takeoff in. Since the legs are so weak (Hubbard 1892; Muir 1906; Linsley & MacSwain 1957), abdomen-catapulting could then increase initial flight speed, and stem-climbing would relax lift requirements. The collisions are unlikely to be caused by poor spatial acuity.

## Applications & Limitations

Recognizing exceptional aerial performance as a pillar of strepsipteran success has great explanatory power. So many other prominent attributes follow on from this apparent necessity, in males: high FMR, low wing loading, prompt flight activity upon adult emergence, reduced body mass, their energetics, and halteres—even strepsipteran vision is optimized for the flight of a tiny (ancestrally nocturnal), pheromone-following insect. To unequivocally demonstrate the centrality of flight to strepsipteran ecology identifying that, e.g., the FMR is high suffices, but precisely how high cannot yet be said; a similar caveat exists for the other aforementioned attributes. Yet the limitations to flight ability have relevance to strepsipteran dispersal, including the inability of adult females to disperse or direct the location of their host. Thus, males must be able to fly well to support expansive female fecundity. If they were unable to do so, adult females would still exit their hosts or Strepsiptera would have evolved to be ordinary parasitoids. These evolutionary decisions, including how Strepsiptera first came to be parasites at all, have profound implications on the evolutionary trajectory of these and, by contrast, other parasitic insects.

## Future directions

Documenting the stages of plume following in a single strepsipteran species or specimen is of great interest. Another important near-future goal is to obtain the actual wet weight of additional Strepsiptera (i.e., besides that of *Stylops ovinae* (Pohl et al., 2020)), along with the wet weight of the thorax, and if feasible, that of the flight muscles themselves. These data would allow for direct calculation of the FMR. Furthermore, the corresponding body length, head and thorax length, thorax width, wing area, and dry weight should also be collected. There should be more research into the developmental state of strepsipteran wings and a careful investigation of whether or not they have a separate wing heart or hearts, or wing circulatory organs at all. Why Strepsiptera have wing buds is also worth considering.

### Laboratory experimentation

When working with adult male Strepsiptera, the advantages of laboratory-based experimentation are tempered by the difficulty of obtaining and transporting material, the general need to keep both parasite and host alive, the brevity of emerged life, and the small size of specimens. Moreover, emerged males have very limited concerns, which are less compatible with known or widely available laboratory techniques. Thus, some natural behaviors fail to present entirely or produce poor results. It can also be difficult to know when reliably repeated behavior is truly representative. Thus, as with noticing Strepsiptera can limit their exposed wing area, sometimes it will be advantageous to perform experiments in the field. Even so, learning how to lab-rear Strepsiptera from additional families is of great importance. (If healthy hosts can be kept alive, stylopized hosts probably can be too.)

#### Specimen immobilization

Refrigeration is often administered to immobilize just-emerged adult males. Refrigeration does not increase their lifespans, however, suggesting that Strepsiptera attempt to compensate for reduced thoracic temperatures in a manner that negatively impacts their energy reserves as potently as active flight does. The physical performance of recently cold-treated specimens may also be altered as they return to normal body temperature.

Another common means of immobilizing insects is to administer CO_2_. Although Strepsiptera cease all movement after only a few seconds of exposure, in my experience, treated specimens never regained the power of flight. In fact, they seem to be impaired in general thereafter, as Kirkpatrick (1937) independently documented, “none so treated recovered properly from any anaesthetic used.”

Alternatively, specimens can be immobilized by physical constraint. It may be most effective to temporarily slip a microscope slide over an emerging specimen and gently reposition it to expose the portion of the body requiring preparation (Ulrich 1933).

#### Avoid interference

Surprisingly, emerged Strepsiptera prevented from flying die within the same time period (2 h) as those allowed to fly (pers. obs.). Thus, emerged adult males can expend as much energy keeping their flight apparatus primed as they do actually flying.

It is unusual for Strepsiptera to become exhausted after only 15 min of flight (Hubbard 1892). However, co-confinement to a tiny area encompassing a nest the resident wasps were obliged to defend, coupled with sex pheromone emanating from multiple calling females into a small volume of air absent any directional wind (David et al. 1983; Kennedy 1983) would provide impetus for exorbitant flight speed maximization, thus consuming a large amount of energy in a very short amount of time.

Friese having observed an emerged *Stylops aterrimus* survive at least 62 h is exceedingly unlikely (Friese 1883; Pierce 1909)—or would be, if the animal had not been removed from its puparium before having reached maturity. Oddly, the account contains nothing concerning the insect’s flight or other activities despite the remarkable span of time they occupied. Friese (1883) writes, ‘despite its forcibly promoted pupal hatching’ [translated from German]; however, its longevity was not *despite*, but *because* of that: Friese did not record the first free-flying Strepsiptera (indirectly indicating that this specimen did not fly) until February 26, about 1.5 months after having actively removed this specimen (Friese 1883). Therefore, Friese’s specimen must have been teneral. It is astoundingly improbable that a Strepsiptera could fly continuously for 2.5 d without feeding—recall that adult Strepsiptera are unable to feed (Pohl & Beutel 2008)—or that such incredible activity would have gone without direct comment.

Whatever Strepsiptera do to prime their flight physiology is apparently inaccessible if their bodies are too teneral. It is incorrect to consider all males confined to pupal cases pharate adults, because they may also be immature. Therefore, one should assume adult male Strepsiptera will liberate themselves at the first best opportunity and normally leave them to it. Caution should be exercised when contemplating accounts of prematurely released Strepsiptera. Strepsipteran self-liberation may facilitate wing elongation and could also constitute a beneficial warmup similar to moths shivering before flying in cold weather. Occasionally, Strepsiptera do not fly immediately after quitting the puparium, even when they initiated and executed the entire procedure autonomously (Hassan 1939; Linsley & MacSwain 1957). Insufficiently warm flight muscles may be a reason why.

## Conclusions

- Strepsiptera are dependent on flight

◦ there are no documented cases of parthenogenesis in Strepsiptera
- Strepsiptera have a well-developed flight apparatus

◦ low-wing loading allows quick turns for following pheromone plumes
◦ high FMR provides force for fighting air currents and quick acceleration
◦ males compete for mates, reinforcing their strong flight apparatus
- Adult males exploit optomotor anemotaxis to track sex pheromone plumes

◦ distinct stages of pheromone tracking account for disparate flight descriptions
- Strepsipteran wings are apparently teneral
- Halteres mark changes in orientation independently of vision

◦ promote high-sensitivity vision by:

▪ allowing spatial acuity and visual-based temporal acuity to degrade with reduced consequence
▪ replacing ocelli (freeing area for greater light gathering)
▪ compensating for lack of polarization sensitivity (freeing area for greater light gathering)
◦ allow rapid response to perturbation
◦ enable better flight control
- Blackberry-esque eyes of adult males are overwhelmingly used to direct flight

◦ ancestral nocturnalism led to increased light sensitivity and decreased spatial acuity
◦ ability to detect difference and direction appear sufficient for optic flow

### Parting shot

The order Strepsiptera (‘twisted wings’) was given its name on account of the deformity of the “forewings” of adult males (i.e., their misidentified “elytra,” which are in fact halteres), and not that of their true wings: “*Strepsiptera* is the term I propose by which to designate the order, which name I have given it on account of its distorted elytra” (Kirby 1813). Thus, the common name, “twisted wing” parasite would be more fitting and informative if rechristened, “crumpled haltere,” which might also accentuate differences in the maturation of the two flight tools—for it appears that the halteres are never teneral.

## Supporting information

Data of body offset supplemental experiment

## Acknowledgments

Much gratitude to SNOMNH for information and lab space. To William Ulch and his mosquito monitoring staff for their by-catch and letting me come on select collection runs. To Dr. Kenneth Hobson and his family and Dr. Scott Russell for accommodation, equipment, and camaraderie, and to my former OK neighbors for all-night access to their backyards. To OU for technical support and lab space. Special thanks to Dr. Peter Follett for providing a stock insect rig. Thank you Kendra and John of Abbott Nature Photography for the vacated spot in their macrophotography workshop. I am grateful to Ursula de Bary, Markus Streicher, and Franziska Leifer for help getting lenses and cameras. Enormous thanks to David Coulon and Ingrid Bush for accommodation as I wrote my dissertation and then this manuscript.

I used the free software packages Docear4Word, JabRef, Irfanview 64, and FastStone Image Viewer. I also made great use of the BHL to obtain historical documents.

## Footnotes

1 It is not clear which *Xenos* sp. Hubbard encountered (but see Brues 1903), or even which *Polistes* (again see Brues (1903)). But, due to its much smaller size and paler coloration, Hubbard (1892) explicitly ruled out *Xenos peckii*.

2 Siphonaptera (fleas) are not parasitic as larvae and are ectoparasites as adults.

3 Presumably, darkness is necessary for the emergence of nocturnal males, but is more readily available than light.

4 Every new arthropod developmental stage (instar) begins while the animal is still enclosed in the husk of its previous instar. This part of the new stage is known as pharate + the name of the new stage (Hinton 1971). Development may continue unabated using the old cuticle as a protective layer. Thus, a pharate adult has separated from its pupal cuticle, but has not cast it off, and may not be mature.

5 That is, the Stylopidia: Strepsiptera in which the female must remain within her host her entire life. In the basal family Mengenillidae, the host reportedly dies shortly after female strepsipteran departure, though it is unclear if the exodus leads directly or indirectly to death.

6 Wing and power loading are most precisely specified in terms of weight, *not* mass. There are high-force flight contexts in which the weight of a flying object would change—affecting flight performance—but its mass would not (Banner 2014). With respect to animals, however, this distinction is unimportant.

7 It is not clear if female Stylopidia have an organ analogous to the male balloon gut. However, at least *Mengenilla moldrzyki* and *Eoxenos laboulbenei* (species of Strepsiptera from Mengenillidae, the most basal extant family for which females are known (Bravo et al. 2009), wherein adult females are facultatively free-living) have a “mouthfield sclerite” (Tröger et al. 2023), which, in male Strepsiptera, functions as an air uptake valve and the opening to the balloon gut (Pohl & Beutel, 2008). There is no chance these wingless females will ever fly, but it would be very useful for them to know when males are likely to do so. Also, in his Fig. 12 in reference to *Corioxenos antestiae* (family Corioxenidae), Cooper (1938) depicts mature unfertilized females as having a large midgut (mesenteron); however, there is no indication of an uptake valve.

8 Once for *Dialeurodes citri* (Byrne et al., 1988); and ostensibly once for *Vespula germanica* (Magnan (1934) (probably a drone) vs. Sotavalta (1952) (presumably a gyne) [I did not have direct access to this book]), both reported in Byrne et al. (1988). However, neither *V. germanica* entry matched well with those from Tercel et al (2018) (most likely workers), so I kept three entries for *V. germanica*.

9 The just noticeable difference (JND)—i.e., the smallest change that can be distinguished 50% of the time under simultaneous viewing conditions—in area is 11.4% (Odic et al. 2013), so flight muscles considered to be extensive (Pohl and Beutel 2008) must be larger than that. [Similar arguments apply to volumetric spaces.]

10 However, I disagree with Pohl and Beutel (2005, 2008) who claimed this was also true of the adult *eyes* of stem group males. Judging from beetle eyes, *Protoxenos* have normal compound eyes with a reduced number of facets: The vertex is broad and the lenses do not protrude beyond it. Thus, the eyes are not abnormally large.

11 Useless to adult male Strepsiptera, which have a high base metabolism upon emergence.

† The dagger symbol indicates an extinct phylogenetic group. Furthermore, in this case, all three are also *stem* groups: they consist entirely of extinct organisms that display more of the morphological features of extant Strepsiptera than any other extant organisms. Thus, these extinct lineages had essentially the same flight apparatus as modern Strepsiptera, but evolved before the most recent common ancestor of all extant strepsipteran taxa.

