## Supplemental Info 1 for "Insights into strepsipteran flight"

### Combined Byrne (1988) and Tercel (2018) data set

| | |
|---|---|
| Paper title: | Insights into strepsipteran flight |
| Jounral name: | *The Science of Nature* |
| Author: | Marisano James |
| Affiliation: | University of California, Davis |
| Email: | |
| Description: | A data set of insect wing loadings compiled from Byrne at al. (1988) and Tercel et al. (2018). The set contains 234 entries, representing 231 species from 11 insect orders, although 4 included orders (Blattodea, Ephemeroptera, Trichoptera, and Mecoptera) each contain a single species. Also includes wingbeat frequencies and notes referencing the original citation. These data are used to evaluate strepsipteran wing loading. As of 2025.12.21, species names have been updated via GBIF to reflect current taxonomy and nomenclature. Spelling and other errors have also been corrected (see below). |

| # | Order | Family | Genus | Species | Mass (g) | Wing area (cm$^2$) | Wing loading (g/cm$^2$) | Wingbeat (Hz) | Average wing loading for Order (g/cm$^2$) | Citation |
|---|---|---|---|---|---|---|---|---|---|---|
| 1 | Hemiptera | Aleyrodidae | *Dialeurodes* | *Dialeurodes citri* (male) | 0.000036 | 0.0207 | 0.00174 | – | 0.03220 | Byrne et al., 1988 |
| 2 | Hemiptera | Aleyrodidae | *Trialeurodes* | *Trialeurodes vaporariorum* | 0.000035 | 0.0165 | 0.00212 | 180.0 | 0.03220 | Byrne et al., 1988 |
| 3 | Hemiptera | Aleyrodidae | *Bemisia* | *Bemisia tabaci* | 0.000033 | 0.0134 | 0.00245 | 168.6 | 0.03220 | Byrne et al., 1988 |
| 4 | Neuroptera | Chrysopidae | *Chrysoperla* | *Chrysoperla carnea* | 0.0065 | 1.972 | 0.003 | 25.923 | 0.0051667 | Tercel et al., 2018 |
| 5 | Neuroptera | Hemerobiidae | *Micromus* | - | 0.0003 | 0.106 | 0.003 | 94.413 | 0.0051667 | Tercel et al., 2018 |
| 6 | Hemiptera | Aleyrodidae | *Dialeurodes* | *Dialeurodes citri* (female) | 0.000080 | 0.0264 | 0.00303 | – | 0.03220 | Byrne et al., 1988 |
| 7 | Hemiptera | Aleyrodidae | *Aleurothrixus* | *Aleurothrixus floccosus* | 0.000065 | 0.194 | 0.00336 | 165.6 | 0.03220 | Byrne et al., 1988 |
| 8 | Lepidoptera | Pieridae | *Pieris* | *Pieris napi* | 0.037 | 8.530 | 0.004 | 6 | 0.06651 | Sotavalta, 1952 |
| 9 | Lepidoptera | Geometridae | *Xanthorhoe* | *Xanthorhoe montanata* | 0.0133 | 2.98 | 0.004 | 29.243 | 0.06651 | Tercel et al., 2018 |
| 10 | Lepidoptera | Nymphalidae | *Aphantopus* | *Aphantopus hyperantus* | 0.0373 | 7.262 | 0.005 | 16.014 | 0.06651 | Tercel et al., 2018 |
| 11 | Hemiptera | Aleyrodidae | *Trialeurodes* | *Trialeurodes abutilonea* | 0.000050 | 0.0096 | 0.00523 | 224.2 | 0.03220 | Byrne et al., 1988 |
| 12 | Diptera | Trichoceridae | *Trichocera* | - (2) | 0.0012 | 0.200 | 0.00600 | 74 | 0.06765 | Sotavalta, 1952 |
| 13 | Lepidoptera | Geometridae | *Pseudopanthera* | *Pseudopanthera macularia* | 0.021 | 3.400 | 0.006 | 25 | 0.06651 | Magnan, 1934 |
| 14 | Neuroptera | Hemerobiidae | *Wesmaelius* | *Wesmaelius subnebulosis* | 0.0026 | 0.416 | 0.006 | 45.304 | 0.0051667 | Tercel et al., 2018 |
| 15 | Neuroptera | Hemerobiidae | *Hemerobius* | *Hemerobius humulinus* | 0.0021 | 0.336 | 0.006 | 46.583 | 0.0051667 | Tercel et al., 2018 |
| 16 | Neuroptera | Hemerobiidae | *Micromus* | *Micromus angulatus* | 0.0059 | 0.932 | 0.006 | 50 | 0.0051667 | Tercel et al., 2018 |
| 17 | Hemiptera | Aphididae | *Acyrthosiphon* | *Acyrthosiphon kondoi* | 0.000702 | 0.1106 | 0.00633 | 81.1 | 0.03220 | Byrne et al., 1988 |
| 18 | Lepidoptera | Pieridae | *Pieris* | *Pieris brassicae* (3) | 0.113 | 15.531 | 0.007 | 11.656 | 0.06651 | Tercel et al., 2018; Magnan, 1934; Sotavalta, 1952 |
| 19 | Neuroptera | Hemerobiidae | *Wesmaelius* | - | 0.0033 | 0.444 | 0.007 | 54.583 | 0.0051667 | Tercel et al., 2018 |
| 20 | Hemiptera | Aphididae | *Aphis* | *Aphis nerii* | 0.000467 | 0.0663 | 0.00750 | 118.1 | 0.03220 | Byrne et al., 1988 |
| 21 | Hemiptera | Aphididae | *Aphis* | *Aphis fabae* | 0.000411 | 0.0526 | 0.00780 | 104.7 | 0.03220 | Byrne et al., 1988 |
| 22 | Lepidoptera | Papilionidae | *Iphiclides* | *Iphiclides podalirius* | 0.300 | 36.000 | 0.008 | 10 | 0.06651 | Magnan, 1934 |
| 23 | Ephemeroptera | Baetidae | *Centroptilum* | *Centroptilum luteolum* | 0.0027 | 0.306 | 0.009 | 75.045 | 0.009 | Tercel et al., 2018 |
| 24 | Hemiptera | Miridae | - | - | 0.0011 | 0.122 | 0.009 | 127.872 | 0.03220 | Tercel et al., 2018 |
| 25 | Lepidoptera | Geometridae | *Pasiphila* | *Pasiphila rectangulata* | 0.0107 | 1.132 | 0.009 | 41.358 | 0.06651 | Tercel et al., 2018 |
| 26 | Lepidoptera | Tortricidae | - | - | 0.0055 | 0.612 | 0.009 | 52.214 | 0.06651 | Tercel et al., 2018 |
| 27 | Lepidoptera | Pieridae | *Gonepteryx* | *Gonepteryx rhamni* | 0.107 | 12.00 | 0.0099 | 21 | 0.06651 | Magnan, 1934 |
| 28 | Lepidoptera | Nymphalidae | *Coenonympha* | *Coenonympha pamphilus* | 0.046 | 4.80 | 0.010 | 22 | 0.06651 | Magnan, 1934 |

| # | Order | Family | Genus | Species | Mass (g) | Wing area ($cm^2$) | Wing loading ($g/cm^2$) | Wingbeat (Hz) | Average wing loading for Order ($g/cm^2$) | Citation |
|---|---|---|---|---|---|---|---|---|---|---|
| 29 | Lepidoptera | Pterophoridae | *Pterophorus* | *Pterophorus pentadactyla* | 0.0114 | 1.12 | 0.01 | 32.333 | 0.06651 | Tercel et al., 2018 |
| 30 | Lepidoptera | Erebidae | *Nyea* | *Nyea lurideola* | 0.0258 | 2.542 | 0.01 | 33.095 | 0.06651 | Tercel et al., 2018 |
| 31 | Lepidoptera | Crambidae | - | - | 0.0059 | 0.562 | 0.01 | 57.948 | 0.06651 | Tercel et al., 2018 |
| 32 | Lepidoptera | Geometridae | *Geometra* | *Geometra papilionaria* | 0.1071 | 10.194 | 0.011 | 22.023 | 0.06651 | Tercel et al., 2018 |
| 33 | Odonata | Calopterygidae | *Calopteryx* | *Calopteryx splendens* | 0.1092 | 9.76 | 0.011 | 19.318 | 0.02854 | Tercel et al., 2018 |
| 34 | Hemiptera | Aphididae | *Aphis* | *Aphis gossypii* | 0.000114 | 0.0103 | 0.01106 | 123.4 | 0.03220 | Byrne et al., 1988 |
| 35 | Diptera | Tipulidae | - | - | 0.002 | 0.168 | 0.012 | 94.606 | 0.06765 | Tercel et al., 2018 |
| 36 | Hemiptera | - | - | - | 0.0014 | 0.114 | 0.012 | 152.247 | 0.03220 | Tercel et al., 2018 |
| 37 | Lepidoptera | Saturniidae | *Samia* | *Samia cynthia* | 0.605 | 50.000 | 0.012 | 8 | 0.06651 | Magnan, 1934 |
| 38 | Lepidoptera | Nymphalidae | *Vanessa* | *Vanessa alalanta* | 0.134 | 10.800 | 0.012 | 10 | 0.06651 | Magnan, 1934 |
| 39 | Lepidoptera | Crambidae | *Anania* | *Anania hortulata* | 0.0293 | 2.41 | 0.012 | 40.996 | 0.06651 | Tercel et al., 2018 |
| 40 | Diptera | Psychodidae | - | - | 0.0006 | 0.048 | 0.013 | 144.611 | 0.06765 | Tercel et al., 2018 |
| 41 | Hemiptera | Aphididae | *Uroleucon* | *Uroleucon cirsii* | 0.0015 | 0.112 | 0.013 | 99.603 | 0.03220 | Tercel et al., 2018 |
| 42 | Lepidoptera | Erebidae | *Hypena* | *Hypena proboscidalis* | 0.0565 | 4.496 | 0.013 | 30.587 | 0.06651 | Tercel et al., 2018 |
| 43 | Lepidoptera | Geometridae | *Idaea* | *Idaea aversata* | 0.0303 | 2.42 | 0.013 | 32.088 | 0.06651 | Tercel et al., 2018 |
| 44 | Lepidoptera | Tortricidae | *Pandemis* | *Pandemis cerasana* | 0.0124 | 0.89 | 0.014 | 54.184 | 0.06651 | Tercel et al., 2018 |
| 45 | Lepidoptera | Tortricidae | *Pseudargyrotoza* | *Pseudargyrotoza conwagana* | 0.0044 | 0.318 | 0.014 | 64.246 | 0.06651 | Tercel et al., 2018 |
| 46 | Lepidoptera | Tortricidae | *Lobesia* | *Lobesia fuligana* | 0.0076 | 0.526 | 0.014 | 64.566 | 0.06651 | Tercel et al., 2018 |
| 47 | Odonata | Calopterygidae | *Calopteryx* | *Calopteryx virgo* | 0.1457 | 10.466 | 0.014 | 17.847 | 0.02854 | Tercel et al., 2018 |
| 48 | Hemiptera | Aphididae | *Myzus* | *Myzus persicae* | 0.000334 | 0.0237 | 0.01412 | 90.9 | 0.03220 | Byrne et al., 1988 |
| 49 | Lepidoptera | Nymphalidae | *Aglais* | *Aglais io* | 0.195 | 14.000 | 0.0147 | 18 | 0.06651 | Magnan, 1934 |
| 50 | Lepidoptera | Nymphalidae | *Argynnis* | *Argynnis pandora* | 0.278 | 18.00 | 0.015 | 10 | 0.06651 | Magnan, 1934 |
| 51 | Lepidoptera | Nymphalidae | *Polygonia* | *Polygonia c-album* | 0.1145 | 7.696 | 0.015 | 27.501 | 0.06651 | Tercel et al., 2018 |
| 52 | Odonata | Coenagrionidae | *Coenagrion* | *Coenagrion puella* (3) | 0.0322 | 2.075 | 0.015 | 37.008 | 0.02854 | Tercel et al., 2018 |
| 53 | Lepidoptera | Saturniidae | *Saturnia* | *Saturnia pyri* | 1.890 | 120.000 | 0.016 | 8 | 0.06651 | Magnan, 1934 |
| 54 | Odonata | Libellulidae | *Perithemis* | *Perithemis tenera* | 0.061 | 3.812 | 0.016 | 39.4 | 0.02854 | May, 1981 |
| 55 | Lepidoptera | Nymphalidae | *Vanessa* | *Vanessa cardui* | 0.173 | 10.40 | 0.017 | 20 | 0.06651 | Magnan, 1934 |
| 56 | Diptera | - | - | - | 0.0009 | 0.05 | 0.018 | 204.355 | 0.06765 | Tercel et al., 2018 |
| 57 | Hemiptera | Miridae | - | - | 0.0048 | 0.25 | 0.019 | 120.832 | 0.03220 | Tercel et al., 2018 |
| 58 | Odonata | Libellulidae | *Libellula* | *Libellula depressa* | 0.245 | 13.200 | 0.019 | 20 | 0.02854 | Magnan, 1934 |
| 59 | Lepidoptera | Erebidae | *Lymantria* | *Lymantria dispar* | 0.101 | 5.39 | 0.02 | 27 | 0.06651 | Casey, 1981 |
| 60 | Odonata | Libellulidae | *Pantala* | *Pantala flavescens* | 0.308 | 15.400 | 0.020 | 22.9 | 0.02854 | May, 1981 |
| 61 | Odonata | Libellulidae | *Tramea* | *Tramea carolina* | 0.358 | 17.900 | 0.020 | 23.6 | 0.02854 | May, 1981 |
| 62 | Hemiptera | - | - | - | 0.0104 | 0.49 | 0.021 | 116.865 | 0.03220 | Tercel et al., 2018 |
| 63 | Hymenoptera | Ichneumonidae | *Ophion* | *Ophion luteus* | 0.033 | 1.5501 | 0.021 | 62 | 0.17692 | Sotavalta, 1952 |
| 64 | Lepidoptera | Erebidae | *Epicallia* | *Epicallia villica* | 0.165 | 8.000 | 0.021 | 20 | 0.06651 | Magnan, 1934 |
| 65 | Odonata | Libellulidae | *Pachydiplax* | *Pachydiplax longipennis* | 0.178 | 8.476 | 0.021 | 24.3 | 0.02854 | May, 1981 |
| 66 | Odonata | Libellulidae | *Tramea* | *Tramea lacerata* | 0.382 | 18.190 | 0.021 | 26.8 | 0.02854 | May, 1981 |
| 67 | Odonata | Libellulidae | *Erythemis* | *Erythemis simplicicollis* | 0.176 | 8.38 | 0.021 | 28 | 0.02854 | May, 1981 |

| # | Order | Family | Genus | Species | Mass (g) | Wing area ($cm^2$) | Wing loading ($g/cm^2$) | Wingbeat (Hz) | Average wing loading for Order ($g/cm^2$) | Citation |
|---|---|---|---|---|---|---|---|---|---|---|
| 68 | Diptera | Chironomidae | - | - (2) | 0.0015 | 0.067 | 0.0215 | 550.92 | 0.06765 | Tercel et al., 2018 |
| 69 | Hymenoptera | Braconidae | - | - | 0.0029 | 0.134 | 0.022 | 136.261 | 0.17692 | Tercel et al., 2018 |
| 70 | Mecoptera | Panorpidae | *Panorpa* | *Panorpa communis* (2) | 0.035 | 1.621 | 0.022 | 38.443 | 0.022 | Magnan, 1934; Tercel et al., 2018 |
| 71 | Odonata | Corduliidae | *Epitheca* | *Epitheca cynosura* | 0.165 | 7.500 | 0.022 | 27.6 | 0.02854 | May, 1981 |
| 72 | Diptera | Syrphidae | *Syrphus* | - | 0.0005 | 0.022 | 0.023 | 190.86 | 0.06765 | Tercel et al., 2018 |
| 73 | Odonata | Libellulidae | *Libellula* | *Libellula luctuosa* | 0.318 | 13.826 | 0.023 | 19 | 0.02854 | May, 1981 |
| 74 | Odonata | Libellulidae | *Orthetrum* | *Orthetrum coerulescens* | 0.248 | 10.800 | 0.023 | 20 | 0.02854 | Magnan, 1934 |
| 75 | Odonata | Libellulidae | *Sympetrum* | *Sympetrum striolatum* | 0.1595 | 6.944 | 0.023 | 40.665 | 0.02854 | Tercel et al., 2018 |
| 76 | Hymenoptera | Torymidae | *Torymus* | - | 0.0024 | 0.098 | 0.024 | 160.011 | 0.17692 | Tercel et al., 2018 |
| 77 | Odonata | Aeshnidae | *Nasiaeschua* | *Nasiaeschua pentacantha* | 0.430 | 16.538 | 0.026 | 21.1 | 0.02854 | May, 1981 |
| 78 | Diptera | Tipulidae | *Tipula* | - (12) | 0.030 | 1.110 | 0.027 | 52 | 0.06765 | Sotavalta, 1952 |
| 79 | Odonata | Libellulidae | *Sympetrum* | *Sympetrum meridionale* | 0.281 | 10.000 | 0.028 | 21 | 0.02854 | Magnan, 1934 |
| 80 | Diptera | Tipulidae | *Nephrotoma* | *Nephrotoma flavescens* | 0.0118 | 0.402 | 0.029 | 79.47 | 0.06765 | Tercel et al., 2018 |
| 81 | Odonata | Libellulidae | *Libellula* | *Libellula quadrimaculatum* (4) | 0.307 | 10.600 | 0.029 | 21 | 0.02854 | Magnan, 1934 |
| 82 | Odonata | Corduliidae | *Somatochlora* | *Somatochlora tenebrosa* | 0.352 | 12.138 | 0.029 | 25.8 | 0.02854 | May, 1981 |
| 83 | Diptera | Tipulidae | *Tipula* | *Tipula maxima* | 0.069 | 2.260 | 0.030 | 48 | 0.06765 | Magnan, 1934 |
| 84 | Hemiptera | Miridae | - | - | 0.0119 | 0.402 | 0.03 | 108.171 | 0.03220 | Tercel et al., 2018 |
| 85 | Lepidoptera | Erebidae | *Calliteara* | *Calliteara pudibunda* | 0.237 | 8.000 | 0.030 | 28 | 0.06651 | Magnan, 1934 |
| 86 | Lepidoptera | Zygaenidae | *Zygaena* | - | 0.0804 | 2.64 | 0.03 | 60.595 | 0.06651 | Tercel et al., 2018 |
| 87 | Odonata | Aeshnidae | *Aeshna* | *Aeshna isoceles* | 0.611 | 17.800 | 0.03 | 20 | 0.02854 | Magnan, 1934 |
| 88 | Odonata | Libellulidae | *Orthetrum* | *Orthetrum cancellatum* | 0.4176 | 13.96 | 0.03 | 38.577 | 0.02854 | Tercel et al., 2018 |
| 89 | Diptera | Culicidae | *Culex* | *Culex pipiens* | 0.0049 | 0.158 | 0.031 | 334.037 | 0.06765 | Tercel et al., 2018 |
| 90 | Hemiptera | Miridae | *Macrolophus* | - | 0.0076 | 0.244 | 0.031 | 139.717 | 0.03220 | Tercel et al., 2018 |
| 91 | Odonata | Libellulidae | *Libellula* | *Libellula pulchella* | 0.508 | 16.387 | 0.031 | 26.2 | 0.02854 | May, 1981 |
| 92 | Coleoptera | Chrysomelidae | *Oulema* | *Oulema melanopus* | 0.0061 | 0.19 | 0.032 | 123.398 | 0.13409 | Tercel et al., 2018 |
| 93 | Hemiptera | - | - | - | 0.0383 | 1.164 | 0.033 | 90.222 | 0.03220 | Tercel et al., 2018 |
| 94 | Lepidoptera | Noctuidae | *Plusia* | *Plusia gamma* | 0.144 | 4.400 | 0.033 | 48 | 0.06651 | Magnan, 1934 |
| 95 | Odonata | Gomphidae | *Ophiogomphus* | *Ophiogomphus cecilia* | 0.312 | 9.400 | 0.033 | 42 | 0.02854 | Magnan, 1934 |
| 96 | Odonata | Libellulidae | *Plathemis* | *Plathemis lydia* | 0.365 | 10.735 | 0.034 | 31.4 | 0.02854 | May, 1981 |
| 97 | Trichoptera | Phryganeidae | *Phryganea* | *Phryganea grandis* | 0.159 | 4.738 | 0.034 | 27.515 | 0.034 | Tercel et al., 2018 |
| 98 | Diptera | Drosophilidae | *Drosophila* | *Drosophila virilis* | 0.0020 | 0.0576 | 0.035[†] | 195 | 0.06765 | Vogel, 1966 |
| 99 | Diptera | Syrphidae | - | - | 0.0044 | 0.124 | 0.035 | 198.89 | 0.06765 | Tercel et al., 2018 |
| 100 | Lepidoptera | Lasiocampidae | *Poecilocampa* | *Poecilocampa populi* | 0.112 | 3.170 | 0.035 | 55 | 0.06651 | Sotavalta, 1952 |
| 101 | Diptera | Culicidae | *Aedes* | *Aedes cantans* | 0.0066 | 0.182 | 0.036 | 286.949 | 0.06765 | Tercel et al., 2018 |
| 102 | Hemiptera | - | - | - | 0.0184 | 0.518 | 0.036 | 112.917 | 0.03220 | Tercel et al., 2018 |
| 103 | Diptera | Chloropidae | *Thaumatomyia* | *Thaumatomyia notata* | 0.0014 | 0.038 | 0.037 | 269.741 | 0.06765 | Tercel et al., 2018 |
| 104 | Hymenoptera | Tenthredinidae | *Athalia* | *Athalia scuttelariae* | 0.0132 | 0.352 | 0.038 | 87.129 | 0.17692 | Tercel et al., 2018 |
| 105 | Lepidoptera | Noctuidae | *Orthosia* | *Orthosia gothica* | 0.1253 | 3.26 | 0.038 | 47.053 | 0.06651 | Tercel et al., 2018 |
| 106 | Diptera | Tipulidae | *Nephrotoma* | *Nephrotoma quadrifaria* | 0.0181 | 0.464 | 0.039 | 67.36 | 0.06765 | Tercel et al., 2018 |

| # | Order | Family | Genus | Species | Mass (g) | Wing area ($cm^2$) | Wing loading ($g/cm^2$) | Wingbeat (Hz) | Average wing loading for Order ($g/cm^2$) | Citation |
|---|---|---|---|---|---|---|---|---|---|---|
| 107 | Diptera | Culicidae | - | - | 0.0058 | 0.150 | 0.039 | 277 | 0.06765 | Sotavalta, 1952 |
| 108 | Diptera | Culicidae | *Aedes* | *Aedes aegypti* (2) | 0.0015 | 0.037 | 0.039 | 480 | 0.06765 | Sotavalta, 1952 |
| 109 | Lepidoptera | Saturniidae | *Automeris* | *Automeris jacunda* (4) | 0.298 | 7.572 | 0.039 | 17.1 | 0.06651 | Bartholomew & Casey, 1978 |
| 110 | Odonata | Aeshnidae | *Anax* | *Anax junius* | 0.820 | 21.02 | 0.039 | 20.5 | 0.02854 | May, 1981 |
| 111 | Odonata | Macromiidae | *Macromia* | *Macromia illinoiensis* | 0.545 | 13.974 | 0.039 | 30.0 | 0.02854 | May, 1981 |
| 112 | Diptera | - | - | - | 0.0039 | 0.098 | 0.04 | 195.996 | 0.06765 | Tercel et al., 2018 |
| 113 | Lepidoptera | Notodontidae | *Pheosia* | *Pheosia tremula* | 0.201 | 5.000 | 0.040 | 22 | 0.06651 | Magnan, 1934 |
| 114 | Lepidoptera | Noctuidae | *Agrotis* | *Agrostis exclamationis* | 0.133 | 3.20 | 0.04 | 41 | 0.06651 | Magnan, 1934 |
| 115 | Lepidoptera | Lasiocampidae | *Malacosoma* | *Malacosoma americanum* | 0.088 | 2.36 | 0.04 | 58 | 0.06651 | Casey, 1981 |
| 116 | Odonata | Aeshnidae | *Aeshna* | *Aeshna mixta* | 0.530 | 13.80 | 0.04 | 38 | 0.02854 | Magnan, 1934 |
| 117 | Diptera | Syrphidae | *Epistrophe* | *Epistrophe gryssulariae* | 0.0200 | 0.480 | 0.042 | 114 | 0.06765 | Weis-Fogh, 1973 |
| 118 | Lepidoptera | Zygaenidae | *Zygaena* | *Zygaena filipendulae* | 0.127 | 3.000 | 0.042 | 48 | 0.06651 | Magnan, 1934 |
| 119 | Diptera | Chloropidae | - | - | 0.003 | 0.07 | 0.043 | 180.05 | 0.06765 | Tercel et al., 2018 |
| 120 | Hymenoptera | Ichneumonidae | - | - | 0.0233 | 0.546 | 0.043 | 110.116 | 0.17692 | Tercel et al., 2018 |
| 121 | Coleoptera | Coccinellidae | *Harmonia* | *Harmonia axyridis* | 0.0283 | 0.644 | 0.044 | 79 | 0.13409 | Tercel et al., 2018 |
| 122 | Diptera | Culicidae | *Culiseta* | *Culiseta annulata* (2) | 0.0073 | 0.162 | 0.046 | 337.66 | 0.06765 | Tercel et al., 2018 |
| 123 | Lepidoptera | Bombycidae | *Macrothylacia* | *Macrothylacia rubi* | 0.595 | 13.000 | 0.046 | 18 | 0.06651 | Magnan, 1934 |
| 124 | Odonata | Macromiidae | *Macromia* | *Macromia taeniolata* | 0.930 | 20.217 | 0.046 | 25.5 | 0.02854 | May, 1981 |
| 125 | Odonata | Aeshnidae | *Brachytron* | *Brachytron pratense* | 0.557 | 12.000 | 0.046 | 33 | 0.02854 | Magnan, 1934 |
| 126 | Hemiptera | Miridae | *Lygus* | *Lygus rugulipennis* | 0.014 | 0.298 | 0.047 | 115.183 | 0.03220 | Tercel et al., 2018 |
| 127 | Coleoptera | Cantharidae | *Rhagonycha* | *Rhagonycha fulva* | 0.0183 | 0.38 | 0.048 | 79.712 | 0.13409 | Tercel et al., 2018 |
| 128 | Diptera | Syrphidae | *Episyrphus* | *Episyrphus balteatus* (8) | 0.0240 | 0.490 | 0.0486 | 141.51 | 0.06765 | Weis-Fogh, 1973; Tercel et al., 2018 |
| 129 | Lepidoptera | Sphingidae | *Laothoe* | *Laothoe populi* | 0.8449 | 17.152 | 0.049 | 29.33 | 0.06651 | Tercel et al., 2018 |
| 130 | Coleoptera | Coccinellidae | *Propylea* | *Propylea 14-punctata* | 0.0105 | 0.21 | 0.05 | 102.427 | 0.13409 | Tercel et al., 2018 |
| 131 | Lepidoptera | Saturniidae | *Automeris* | *Automeris zugana* (4) | 0.420 | 8.393 | 0.050 | 18.6 | 0.06651 | Bartholomew & Casey, 1978 |
| 132 | Diptera | Anthomyiidae | *Fannia* | *Fannia scalaris* | 0.010 | 0.196 | 0.051 | 210 | 0.06765 | Magnan, 1934 |
| 133 | Lepidoptera | Saturniidae | *Hyperchirica* | *Hyperchirica nausica* (2) | 0.200 | 3.950 | 0.051 | 21.6 | 0.06651 | Bartholomew & Casey, 1978 |
| 134 | Odonata | Aeshnidae | *Anax* | *Anax imperator* | 1.200 | 22.800 | 0.053 | 22 | 0.02854 | Magnan, 1934 |
| 135 | Odonata | Aeshnidae | *Aeshna* | *Aeshna grandis* | 1.2296 | 22.784 | 0.054 | 31.214 | 0.02854 | Tercel et al., 2018 |
| 136 | Diptera | Syrphidae | *Platycheirus* | *Platycheirus peltatus* | 0.0128 | 0.230 | 0.056 | 147 | 0.06765 | Weis-Fogh, 1973 |
| 137 | Coleoptera | Scarabaeidae | *Aphodius* | - | 0.0327 | 0.56 | 0.058 | 93.054 | 0.13409 | Tercel et al., 2018 |
| 138 | Diptera | Tipulidae | *Tipula* | - | 0.0676 | 1.17 | 0.058 | 59.567 | 0.06765 | Tercel et al., 2018 |
| 139 | Diptera | Stratiomyidae | *Chloromyia* | *Chloromyia formosa* | 0.0183 | 0.312 | 0.059 | 156.043 | 0.06765 | Tercel et al., 2018 |
| 140 | Diptera | Culicidae | *Theobaldia* | *Theobaldia annulata* | 0.0099 | 0.169 | 0.059 | 262 | 0.06765 | Sotavalta, 1952 |
| 141 | Diptera | Muscidae | *Musca* | *Musca domestica* | 0.012 | 0.200 | 0.060 | 190 | 0.06765 | Magnan, 1934 |
| 142 | Diptera | Scathophagidae | *Scathophaga* | *Scathophaga stercoraria* | 0.0224 | 0.366 | 0.061 | 104.015 | 0.06765 | Tercel et al., 2018 |
| 143 | Diptera | Tabanidae | *Haematopota* | *Haematopota pluvialis* | 0.0183 | 0.302 | 0.061 | 151.568 | 0.06765 | Tercel et al., 2018 |
| 144 | Diptera | Syrphidae | *Eupeodes* | *Eupeodes corollae* (3) | 0.0213 | 0.350 | 0.061 | 174 | 0.06765 | Weis-Fogh, 1973 |
| 145 | Lepidoptera | Saturniidae | *Automeris* | *Automeris belti* | 0.665 | 10.960 | 0.061 | 14.4 | 0.06651 | Bartholomew & Casey, 1978 |

| # | Order | Family | Genus | Species | Mass (g) | Wing area (cm$^2$) | Wing loading (g/cm$^2$) | Wingbeat (Hz) | Average wing loading for Order (g/cm$^2$) | Citation |
|---|---|---|---|---|---|---|---|---|---|---|
| 146 | Lepidoptera | Noctuidae | *Noctua* | *Noctua pronuba* | 0.485 | 7.800 | 0.062 | 24 | 0.06651 | Magnan, 1934 |
| 147 | Hymenoptera | Vespidae: Eumeninae | - | - | 0.0184 | 0.292 | 0.063 | 135.597 | 0.17692 | Tercel et al., 2018 |
| 148 | Diptera | Empididae | - | - | 0.0193 | 0.288 | 0.067 | 151.321 | 0.06765 | Tercel et al., 2018 |
| 149 | Lepidoptera | Saturniidae | *Hylesia* | - (5) | 0.168 | 2.334 | 0.072 | 32.4 | 0.06651 | Bartholomew & Casey, 1978 |
| 150 | Lepidoptera | Sphingidae | *Hemaris* | *Hemaris fuciformis* | 0.189 | 2.620 | 0.072 | 80 | 0.06651 | Magnan, 1934 |
| 151 | Diptera | Syrphidae | *Eupeodes* | *Eupeodes nitens* | 0.022 | 0.300 | 0.073 | 172 | 0.06765 | Weis-Fogh, 1973 |
| 152 | Diptera | Syrphidae | *Syrphus* | *Syrphus ribesii* (4) | 0.346 | 0.465 | 0.0748 | 183.977 | 0.06765 | Tercel et al., 2018; Weis-Fogh, 1973 |
| 153 | Lepidoptera | Sphingidae | *Deilephila* | *Deilephila elpenor* | 0.5281 | 6.792 | 0.078 | 53.715 | 0.06651 | Tercel et al., 2018 |
| 154 | Hymenoptera | Braconidae | - | - | 0.003 | 0.038 | 0.079 | 164.443 | 0.17692 | Tercel et al., 2018 |
| 155 | Lepidoptera | Sphingidae | *Macroglossum* | *Macroglossum stellatarum* (2) | 0.314 | 3.895 | 0.080 | 79 | 0.06651 | Magnan, 1934; Sotavalta, 1952 |
| 156 | Diptera | Syrphidae | *Syrphus* | *Syrphus vitripennis* | 0.0385 | 0.480 | 0.080 | 196 | 0.06765 | Weis-Fogh, 1973 |
| 157 | Lepidoptera | Crambidae | *Chrysoteuchia* | *Chrysoteuchia culmella* | 0.109 | 1.33 | 0.082 | 40.626 | 0.06651 | Tercel et al., 2018 |
| 158 | Diptera | Syrphidae | *Scaeva* | *Scaeva pyrastri* | 0.034 | 0.400 | 0.085 | 190 | 0.06765 | Magnan, 1934 |
| 159 | Lepidoptera | Sphingidae | *Acherontia* | *Acherontia atropos* (2) | 1.920 | 21.931 | 0.087 | 25.58 | 0.06651 | Magnan, 1934; Tercel et al., 2018 |
| 160 | Lepidoptera | Saturniidae | *Eacles* | *Eacles imperialis* (3) | 1.105 | 12.600 | 0.088 | 17.9 | 0.06651 | Bartholomew & Casey, 1978 |
| 161 | Lepidoptera | Saturniidae | *Automerina* | *Automerina auletes* | 0.720 | 8.090 | 0.089 | 23.4 | 0.06651 | Bartholomew & Casey, 1978 |
| 162 | Coleoptera | Oedemeridae | *Oedemera* | *Oedemera nobilis* | 0.021 | 0.232 | 0.091 | 112.656 | 0.13409 | Tercel et al., 2018 |
| 163 | Diptera | Calliphoridae | *Calliphora* | *Calliphora vicina* (13) | 0.051 | 0.552 | 0.092 | 162 | 0.06765 | Sotavalta, 1952; Magnan, 1934 |
| 164 | Coleoptera | Cantharidae | *Telephorus* | *Telephorus fuscus* | 0.109 | 1.160 | 0.094 | 72 | 0.13409 | Magnan, 1934 |
| 165 | Diptera | Syrphidae | *Sphaerophoria* | *Sphaerophoria scripta* (3) | 0.0193 | 0.200 | 0.094 | 308 | 0.06765 | Weis-Fogh, 1973 |
| 166 | Diptera | Syrphidae | *Chrysotoxum* | *Chrysotoxum arcuatum* | 0.073 | 0.740 | 0.099 | 144 | 0.06765 | Magnan, 1934 |
| 167 | Hemiptera | Cicadidae | *Cicada* | - | 0.752 | 7.64 | 0.10 | 42 | 0.03220 | Ahmad, 1984 |
| 168 | Diptera | Sarcophagidae | *Sarcophaga* | - | 0.054 | 0.526 | 0.103 | 149.643 | 0.06765 | Tercel et al., 2018 |
| 169 | Lepidoptera | Saturniidae | *Automeris* | *Automeris hamata* | 0.564 | 5.450 | 0.103 | 23.5 | 0.06651 | Bartholomew & Casey, 1978 |
| 170 | Coleoptera | Cerambycidae | - | - | 0.142 | 1.33 | 0.107 | 80 | 0.13409 | Sotavalta, 1952 |
| 171 | Diptera | Syrphidae | *Chrysotoxum* | *Chrysotoxum vernale* | 0.064 | 0.600 | 0.107 | 150 | 0.06765 | Magnan, 1934 |
| 172 | Hymenoptera | Apidae | *Andrena* | - | 0.0376 | 0.352 | 0.107 | 213.815 | 0.17692 | Tercel et al., 2018 |
| 173 | Diptera | Syrphidae | *Chrysotoxum* | *Chrysotoxum bicinctum* | 0.075 | 0.680 | 0.110 | 120 | 0.06765 | Magnan, 1934 |
| 174 | Hymenoptera | Sphecidae | *Ammophila* | *Ammophila sabulosa* | 0.045 | 0.420 | 0.11 | 120 | 0.17692 | Magnan, 1934 |
| 175 | Hymenoptera | Apidae | *Apis* | - | 0.0213 | 0.20 | 0.11 | 130 | 0.17692 | Ahmad, 1984 |
| 176 | Diptera | Syrphidae | - | - | 0.0385 | 0.34 | 0.113 | 208.54 | 0.06765 | Tercel et al., 2018 |
| 177 | Lepidoptera | Sphingidae | *Manduca* | *Manduca lefeburii* | 0.571 | 4.920 | 0.116 | 33.1 | 0.06651 | Bartholomew & Casey, 1978 |
| 178 | Diptera | Syrphidae | *Volucella* | *Volucella pellucens* (2) | 0.117 | 0.966 | 0.117 | 127.09 | 0.06765 | Magnan, 1934; Tercel et al., 2018 |
| 179 | Hemiptera | Pentatomidae | *Pentatoma* | *Pentatoma rufipes* | 0.1397 | 1.186 | 0.118 | 96.667 | 0.03220 | Tercel et al., 2018 |
| 180 | Coleoptera | Cerambicidae | *Leptura* | *Leptura quadrifasciata* | 0.1173 | 0.982 | 0.119 | 93.768 | 0.13409 | Tercel et al., 2018 |
| 181 | Diptera | Calliphoridae | *Calliphora* | *Calliphora vomitoria* | 0.0549 | 0.46 | 0.119 | 214.835 | 0.06765 | Tercel et al., 2018 |
| 182 | Hymenoptera | Vespidae | *Vespula* | *Vespula vulgaris* (3) | 0.0891 | 0.755 | 0.121 | 153 | 0.17692 | Sotavalta, 1952; Tercel et al., 2018 |
| 183 | Diptera | Sarcophagidae | *Sarcophaga* | *Sarcophaga carnaria* | 0.045 | 0.36 | 0.125 | 160 | 0.06765 | Magnan, 1934 |
| 184 | Coleoptera | Melolonthidae | *Amphimallon* | *Amphimallon solstitalis* | 0.291 | 2.290 | 0.127 | 78 | 0.13409 | Sotavalta, 1952 |

| # | Order | Family | Genus | Species | Mass (g) | Wing area ($cm^2$) | Wing loading ($g/cm^2$) | Wingbeat (Hz) | Average wing loading for Order ($g/cm^2$) | Citation |
|---|---|---|---|---|---|---|---|---|---|---|
| 185 | Hymenoptera | Apidae | *Andrena* | - | 0.0453 | 0.346 | 0.131 | 172.581 | 0.17692 | Tercel et al., 2018 |
| 186 | Lepidoptera | Sphingidae | *Enyo* | *Enyo ocypete* | 0.388 | 2.950 | 0.132 | 56.1 | 0.06651 | Bartholomew & Casey, 1978 |
| 187 | Coleoptera | Cerambicidae | *Rutpela* | *Rutpela maculata* | 0.1026 | 0.768 | 0.134 | 86.84 | 0.13409 | Tercel et al., 2018 |
| 188 | Coleoptera | Scarabaeidae | *Aphodius* | - (2) | 0.101 | 0.74 | 0.137 | 102.135 | 0.13409 | Tercel et al., 2018 |
| 189 | Hymenoptera | Vespidae | *Vespula* | *Vespula germanica* (3) | 0.081 | 0.591 | 0.137 | 148 | 0.17692 | Tercel et al., 2018 (most likely workers) |
| 190 | Diptera | Syrphidae | *Eristalis* | *Eristalis tenax* (8) | 0.12 | 0.814 | 0.147 | 182.5 | 0.06765 | Magnan, 1934; Weis-Fogh, 1973; Sotavalta, 1952 |
| 191 | Lepidoptera | Sphingidae | *Pachygonidia* | *Pachygonidia drucei* | 0.702 | 4.770 | 0.147 | 48.4 | 0.06651 | Bartholomew & Casey, 1978 |
| 192 | Hymenoptera | Crabronidae | *Ectemnius* | *Ectemnius cavifrons* | 0.08 | 0.542 | 0.148 | 210.688 | 0.17692 | Tercel et al., 2018 |
| 193 | Blattodea | Blattidae | *Periplanela* | *Periplanela americana* | 1.555 | 10.44 | 0.149 | 26 | 0.149 | Ahmad, 1984 |
| 194 | Diptera | Tabanidae | *Tabanus* | *Tabanus bovinus* | 0.276 | 1.840 | 0.150 | 96 | 0.06765 | Magnan, 1934 |
| 195 | Lepidoptera | Saturniidae | *Adeloneivaia* | *Adeloneivaia boisduvalii* (3) | 0.839 | 5.564 | 0.151 | 24.9 | 0.06651 | Bartholomew & Casey, 1978 |
| 196 | Diptera | Syrphidae | *Volucella* | *Volucella bombylans* (2) | 0.1432 | 0.943 | 0.1515 | 126.539 | 0.06765 | Magnan, 1934; Tercel et al., 2018 |
| 197 | Diptera | Tabanidae | *Dasyrhamphis* | *Dasyrhamphis ater* | 0.233 | 1.500 | 0.155 | 100 | 0.06765 | Magnan, 1934 |
| 198 | Hymenoptera | Apidae | *Apis* | *Apis mellifera* (7) | 0.087 | 0.56 | 0.156 | 234.00 | 0.17692 | Magnan, 1934; Sotavalta, 1952; Tercel et al., 2018 |
| 199 | Lepidoptera | Sphingidae | *Xylophones* | *Xylophones libya* | 0.559 | 3.560 | 0.157 | 48.5 | 0.06651 | Bartholomew & Casey, 1978 |
| 200 | Lepidoptera | Sphingidae | *Manduca* | *Manduca corallina* (4) | 1.618 | 10.270 | 0.158 | 28.0 | 0.06651 | Bartholomew & Casey, 1978 |
| 201 | Lepidoptera | Saturniidae | *Syssphinx* | *Syssphinx molina* | 1.630 | 9.700 | 0.168 | 22.9 | 0.06651 | Bartholomew & Casey, 1978 |
| 202 | Lepidoptera | Saturniidae | *Adeloneivaia* | *Adeloneivaia subungulata* | 0.487 | 2.900 | 0.168 | 41.0 | 0.06651 | Bartholomew & Casey, 1978 |
| 203 | Diptera | Syrphidae | *Eristalis* | *Eristalis arbustorum* | 0.0705 | 0.410 | 0.172 | 211 | 0.06765 | Weis-Fogh, 1973 |
| 204 | Hymenoptera | Apidae | *Bombus* | *Bombus hortorum* | 0.159 | 0.900 | 0.177 | 135 | 0.17692 | Magnan, 1934 |
| 205 | Coleoptera | Scutelleridae | *Chrysocoris* | *Chrysocoris purpureus* | 0.264 | 1.50 | 0.18 | 100 | 0.13409 | Ahmad, 1984 |
| 206 | Hymenoptera | Vespidae | *Vespula* | *Vespula germanica* | 0.240 | 1.330 | 0.180 | 139 | 0.17692 | Sotavalta, 1952 (presumably, a gyne) |
| 207 | Coleoptera | Melolonthidae | *Melolontha* | *Melolontha vulgaris* (2) | 0.779 | 4.235 | 0.187 | 54 | 0.13409 | Magnan, 1934; Sotavalta, 1952 |
| 208 | Lepidoptera | Sphingidae | *Xylophones* | *Xylophones pluto* (2) | 0.829 | 4.430 | 0.187 | 45.0 | 0.06651 | Bartholomew & Casey, 1978 |
| 209 | Hymenoptera | Apidae | *Exaerete* | *Exaerete frontalis* (4) | 0.644 | 3.48 | 0.19 | 87 | 0.17692 | Casey et al., 1985 |
| 210 | Hymenoptera | Apidae | *Bombus* | *Bombus pascuorum* | 0.1166 | 0.614 | 0.19 | 198.274 | 0.17692 | Tercel et al., 2018 |
| 211 | Hymenoptera | Vespidae | *Vespula* | *Vespula germanica* | 0.187 | 0.980 | 0.191 | 110 | 0.17692 | Magnan, 1934 (probably a drone) |
| 212 | Hymenoptera | Apidae | *Bombus* | *Bombus terrestris* (9) | 0.216 | 0.128 | 0.194 | 158.507 | 0.17692 | Magnan, 1934; Tercel et al., 2018 |
| 213 | Hymenoptera | Apidae | *Euglossa* | *Euglossa mandibularis* (4) | 0.090 | 0.44 | 0.20 | 209 | 0.17692 | Casey et al., 1985 |
| 214 | Hymenoptera | Apidae | *Euglossa* | *Euglossa saphirina* | 0.071 | 0.35 | 0.20 | 265 | 0.17692 | Casey et al., 1985 |
| 215 | Hymenoptera | Vespidae | *Vespa* | *Vespa crabro* (2) | 0.582 | 2.820 | 0.207 | 102 | 0.17692 | Magnan, 1934; Sotavalta, 1952 |
| 216 | Hymenoptera | Apidae | *Englossa* | *Englossa imperialis* (6) | 0.169 | 0.79 | 0.21 | 179 | 0.17692 | Casey et al., 1985 |
| 217 | Lepidoptera | Sphingidae | *Erinnyis* | *Erinnyis ello* | 1.210 | 5.480 | 0.221 | 23.7 | 0.06651 | Bartholomew & Casey, 1978 |
| 218 | Hymenoptera | Apidae | *Euglossa* | *Euglossa dissimula* | 0.104 | 0.45 | 0.23 | 220 | 0.17692 | Casey et al., 1985 |
| 219 | Hemiptera | Tessaratomidae | *Tesseratoma* | *Tesseratoma javanica* | 0.926 | 3.88 | 0.239 | 66 | 0.03220 | Ahmad, 1984 |
| 220 | Hymenoptera | Apidae | *Eulaema* | *Eulaema nigrita* (2) | 0.399 | 1.67 | 0.24 | 149 | 0.17692 | Casey et al., 1985 |
| 221 | Hymenoptera | Vespidae | *Polistes* | *Polistes gallicus* | 0.115 | 0.460 | 0.250 | 220 | 0.17692 | Magnan, 1934 |
| 222 | Hymenoptera | Apidae | *Bombus* | *Bombus muscorum* | 0.226 | 0.900 | 0.251 | 128 | 0.17692 | Magnan, 1934 |
| 223 | Lepidoptera | Sphingidae | *Manduca* | *Manduca rustica* | 2.704 | 10.720 | 0.252 | 29.5 | 0.06651 | Bartholomew & Casey, 1978 |

| # | Order | Family | Genus | Species | Mass (g) | Wing area ($cm^2$) | Wing loading ($g/cm^2$) | Wingbeat (Hz) | Average wing loading for Order ($g/cm^2$) | Citation |
|---|---|---|---|---|---|---|---|---|---|---|
| 224 | Lepidoptera | Sphingidae | *Perigonia* | *Perigonia lusca* (2) | 0.638 | 2.470 | 0.258 | 62.9 | 0.06651 | Bartholomew & Casey, 1978 |
| 225 | Hymenoptera | Apidae | *Eulaema* | *Eulaema meriana* (7) | 0.940 | 3.46 | 0.270 | 98 | 0.17692 | Casey et al., 1985 |
| 226 | Hymenoptera | Apidae | *Eulaema* | *Eulaema cingulata* (6) | 0.547 | 2.03 | 0.27 | 128 | 0.17692 | Casey et al., 1985 |
| 227 | Lepidoptera | Sphingidae | *Oryba* | *Oryba achemenides* (2) | 2.809 | 10.200 | 0.275 | 39.9 | 0.06651 | Bartholomew & Casey, 1978 |
| 228 | Hymenoptera | Apidae | *Bombus* | *Bombus lapidarius* (4) | 0.446 | 1.472 | 0.29375 | 148.387 | 0.17692 | Magnan, 1934; Sotavalta, 1952; Tercel et al., 2018 |
| 229 | Coleoptera | Lucanidae | *Lucanus* | *Lucanus cervus* | 2.600 | 8.000 | 0.325 | 33 | 0.13409 | Magnan, 1934 |
| 230 | Hymenoptera | Apidae | *Eufriesia* | *Eufriesia pulchra* (4) | 0.425 | 1.27 | 0.33 | 170 | 0.17692 | Casey et al., 1985 |
| 231 | Hymenoptera | Xylocopidae | *Xylocopa* | *Xylocopa violacea* | 0.614 | 1.720 | 0.357 | 130 | 0.17692 | Magnan, 1934 |
| 232 | Lepidoptera | Sphingidae | *Madoryx* | *Madoryx oiclus* (2) | 1.699 | 4.715 | 0.360 | 41.8 | 0.06651 | Bartholomew & Casey, 1978 |
| 233 | Coleoptera | Cetoniidae | *Cetonia* | *Cetonia aurata* | 0.537 | 1.300 | 0.413 | 86 | 0.13409 | Magnan, 1934 |
| 234 | Hymenoptera | Apidae | *Bombus* | - | 1.600 | 3.50 | 0.46 | 125 | 0.17692 | Ahmad, 1984 |

†Vogel (1966) found a single wing area of 2.88 $mm^2$, which should have resulted in an overall wing area of 0.0576 $cm^2$ for both wings, rather than 0.058 $cm^2$, as reported in Byrne at al. (1988). The rounded value produces a corresponding wing loading of 0.035, rather than the otherwise computed 0.034 $g/cm^2$. I do not know why the value was rounded up; four decimal digits are presented in several other instances. I also found that the mass and wing area data from Weis-Fogh (1972) for this species (*Drosophila virilis*) actually use this same data from Vogel (1966), so I discarded that entry—#14 from Byrne et al. (1988). (The entry also contained an error in the wing loading. Byrne et al. (1988) may have copied a value Weis-Fogh (1972) rounded up, and then rounded it up again, producing a very different result for two rows containing the same initial data for the calculation.)

#### Compiled from:

Byrne DN, Buchmann SL, Spangler HG (1988) Relationship between wing loading, wingbeat frequency and body mass in homopterous insects. J Exp Biol 135: 9–23

Tercel MPTG, Veronesi F, Pope TW (2018) Phylogenetic clustering of wingbeat frequency and flight-associated morphometrics across insect orders. Physiol Entomol 43: 149–157

#### Internal references:

Ahmad A (1984) A comparative study on flight surface and aerodynamic parameters of insects, birds and bats. J Exp Biol 22: 270–278

Bartholomew GA, Case TM (1978) Oxygen consumption of moths during rest, pre-flight warm-up, and flight in relation to body size and wing morphology, J Exp Biol 76: 11–25

Casey TM (1981) A comparison of mechanical and energetic estimates of flight cost for hovering sphinx moths. J Exp Biol 91: 117–129

Casey TM, May ML, Morgan KR (1985) Flight energetics of euglossine bees in relation to morphology and wing stroke frequency. J Exp Biol 39: 271–289

Magnan A. (1934) La locomotion chez les animaux 1: Le vol des insectes. Hermann & Co, Paris

May ML (1981) Wingstroke frequency of dragonflies (Odonata: Anisoptera) in relation of temperature and body size. J Comp Physiol 144: 229–240

Sotavalta O. (1952) The essential factor regulating the wing-stroke frequency of insects in wing mutilation and loading experiments and in experiments at subatmospheric pressure. Annales zoologici Societatis Zoologicae Botanicae Fennicae Vanamo, Helsinki

Vogel S. (1966) Flight in Drosophila: I. Flight performance of tethered flies. J Exp Biol 44: 567–578

Weis-Fogh T (1973) Quick estimates of flight fitness in hovering animals, including novel mechanisms for lift production. J Exp Biol 59: 169–230

**Paper title:** Insights into strepsipteran flight

**Journal name:** *Scientific Reports*

**Author:** Marisano James

**Affiliation:** University of California, Davis

**Description:** Contains additional data for pertaining to wing areas, body length vs. wing area, ventral clap-and-fling, and approximate strepsipteran wing stroke amplitude obtained from suitable single images.

#### *Wing areas (*Triozocera texana*)*

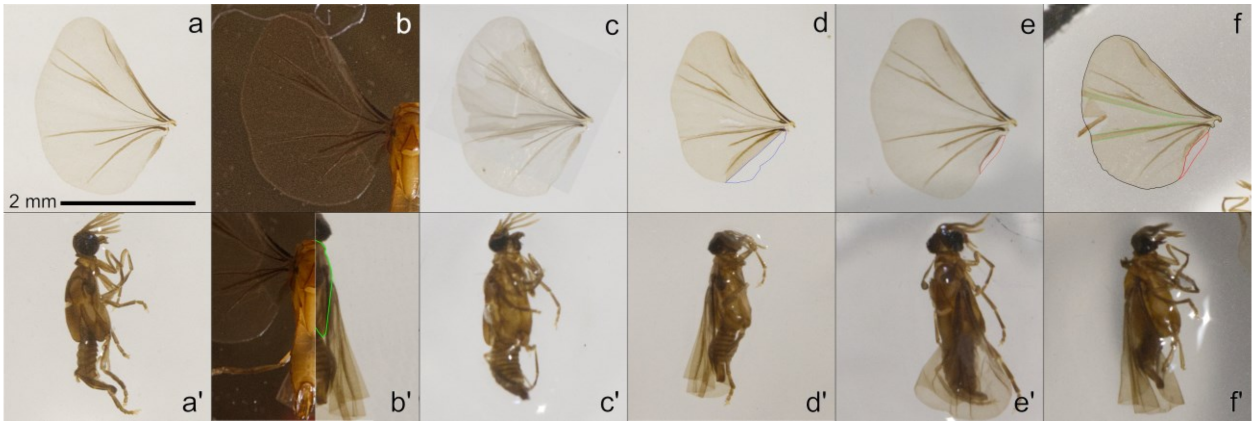


**Fig. S1** Wing areas and body lengths. Six *Triozocera texana* wings and the bodies from which there were removed are shown. The wing-body pairs are presented from least to most augmented. Given how extremely delicate strepsipteran wings are, in all but the first two *(a)* & *(b)*, some digital manipulation was necessary to arrive at a complete wing. However, because in *(b')* the head was removed for analysis before the body length was measured, only *(a + a')* was used to calculate the wing loading. Nonetheless, all of the wings are of similar shape. (a) An entirely detached fully unfurled *T. texana* wing. (b) Outstretched wing measured while still attached to the (headless) body. (b') To avoid abdomen shrinkage issues while approximating the body length of this headless specimen, the thoraces of several headed specimens were resized to fit to that of the target insect, and the resultant average head length was added to that of the headless body. In this image, the headless body appears at left and one of the fitted bodies at right. Note that the ethanol-solution immersed abdomen of the target specimen has shrunk less than that of the dried specimen presented to the right. (c) The left and right wings of this specimen were torn, but because one tore at the bottom and the other at the top, they were fit together to form one complete wing. (d) This wing was fine except that it contains a darkened folded over portion at its base. It has been digitally traced, flipped outward, and its area (blue) added to that of the rest of the wing. (e) The proximal part of the wing base was apparently lost. I fit wing *(a)* to it and added the additional area outlined in red. (f) Many adjustments were made to this wing. Areas darkened due to overlap (outlined in green) were added twice. The ripped portion at the trailing edge of the wing was reconnected to complete the wing profile. Finally, a portion of the base of the wing that had been torn away was traced from another wing, resized and repositioned to closely match the target wing, and added to its base (in red). Note that expanding the green areas would move the wing's leading edge spar into a position similar to that of the other five wings. All images are drawn to scale.

#### *Body length vs. wing area (*Triozocera texana*)*

In the single instance in which it was used to estimate body mass, body length was measured from the base of the antennae (i.e., the outer edge of the eye) to the tip of the aedeagus. In general, however, body lengths of adult male Strepsiptera are unreliable due to abdominal shrinkage subsequent to death. Shrinkage is most pronounced in dried specimens, but is often present in ethanol-preserved specimens, as well. Therefore, to compare wing length to wing area in Fig. S2, only the head and thorax were used to compute “body” length. The resultant curve fits are very strong, indicating that the wing area calculations involving folded over areas, small peripheral tears, and reconstructions from partial wings are very likely to be highly accurate, although those data were not actually used.

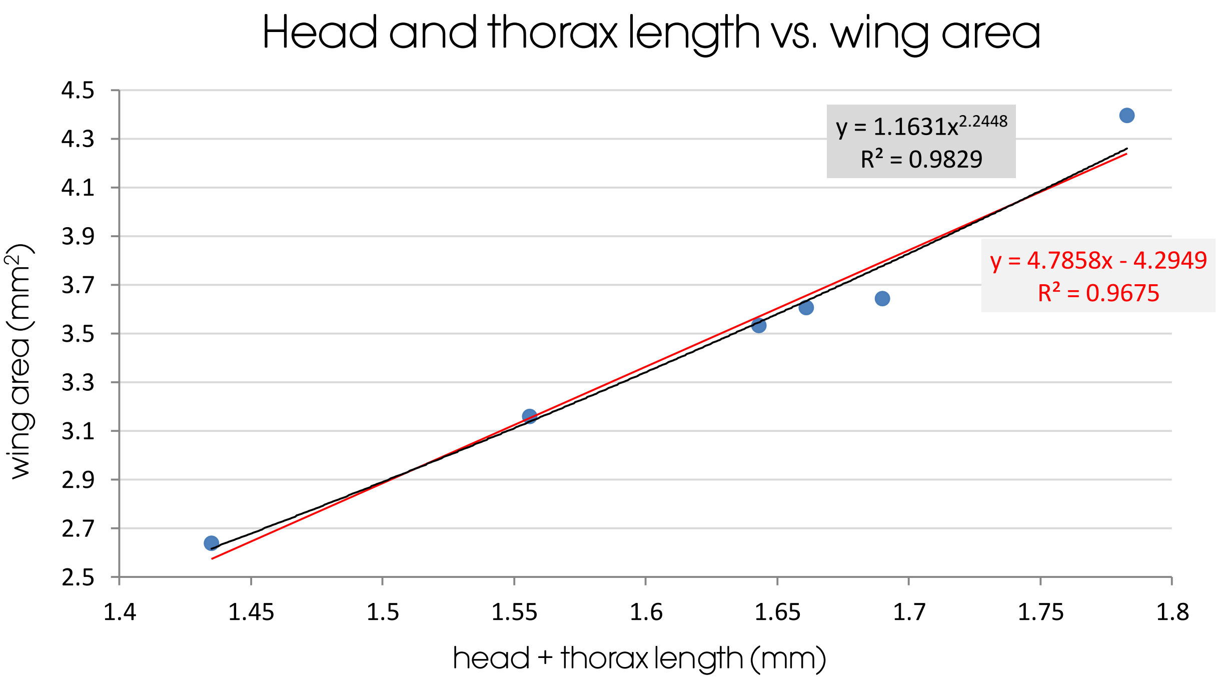


**Fig. S2** Head and thorax length vs. wing area. Here, the head and thorax length of the six *Triozocera texana* depicted in Fig. S1 have been plotted against their wing areas. A nearly linear relationship ($R^2 \approx 0.97$) between wing area and head + thorax length was found, which should also hold for the entire body length, provided the abdomens are relaxed (i.e., neither shrunken nor distended). Fitting to a (super quadratic) power curve produces a slightly better result ($R^2 \approx 0.98$). Therefore, wing area appears to vary systematically with body length (and weight) in *T. texana*, and by extension, in Strepsiptera at large.

#### *Ventral clap-and-fling*

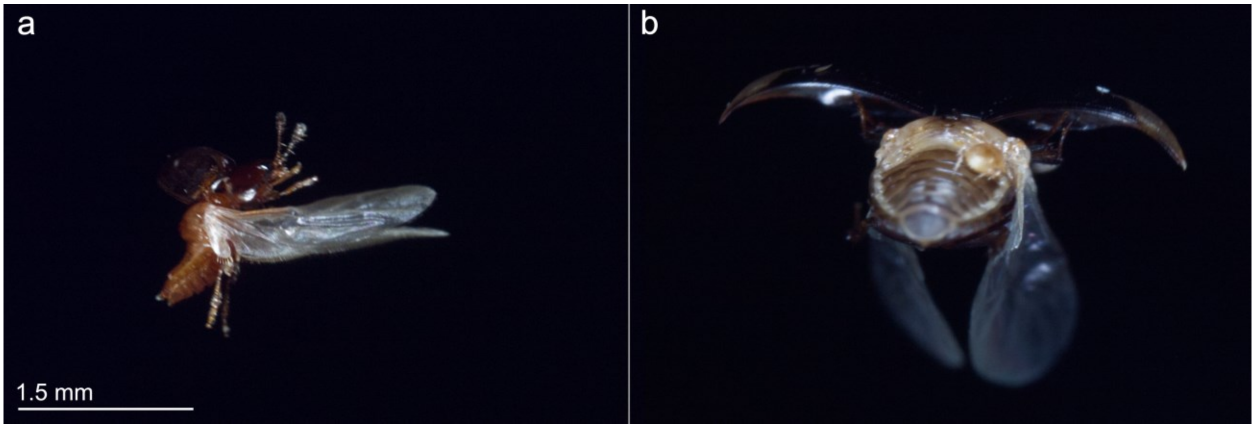


**Fig. S3** Ventral clap-and-fling. As mentioned in the body of the paper, the seven-spotted ladybug (*Coccinella septempunctata*) has been shown to exercise the lift enhancing clap-and-fling mechanism at the end of its downstroke (Yang et al. 2024) in addition to the end of its upstroke, as is typical for the clap-and-fling variants. The use of clap-and-fling at the end of the downstroke had previously been undocumented. To that, I add two instances of ventral lift enhancement in free-flying non-coccinellid beetles that I documented using a heavily modified commercial insect rig in what was my backyard in Oklahoma. Unlike what Yang et al. (2024) documented, neither of these occurred during climbing flight: (a) A tiny beetle employing clap-and-peel at the end of its downstroke. Judging from the thoracic bulge at the base of its wings, this insect has a high FMR. (b) A third beetle species employing a ventral clap-and-fling variant, most likely a near clap-and-fling. (Also, note the mite on its abdomen.) Both images were taken on the evening of 22 VIII 2021. Thus, at least several smaller beetles can further enhance lift via ventral clap-and-fling, which may allow them to fly with even higher power loading (i.e., with even less powerful flight musclature), although a *(a)* strongly suggests, this need not be the case.

#### *Wingstroke amplitude*

Strepsipteran halteres beat in antiphase to their wings (Pix et al. 1993), so the wingstroke amplitude can potentially be estimated from single still images when the body axis is agreeable and the halteres are extended directly above the thorax.[1] Two photographs were captured wherein both criteria were fulfilled for *Triozocera. texana* in free flight. From them, I measured the angular offset from the central axis of the juxtaposed halteres to the leading edge of the outwardly opened wing (Fig. S4). The larger the angular extent of a wingstroke and the faster the wings flap, the more force is produced and the greater the energy consumption. Greater force produces greater speed, but can also be used to takeoff more readily or to hover. I estimated the *T. texana* wingstroke amplitude to be about 154°, using the average measurement taken from those two photographs of separate free-flying specimens (Fig. S4). Pix et al. (1993) measured the wingstroke amplitude of *Xenos vesparum* to be about 160° using conventional high-speed videography.

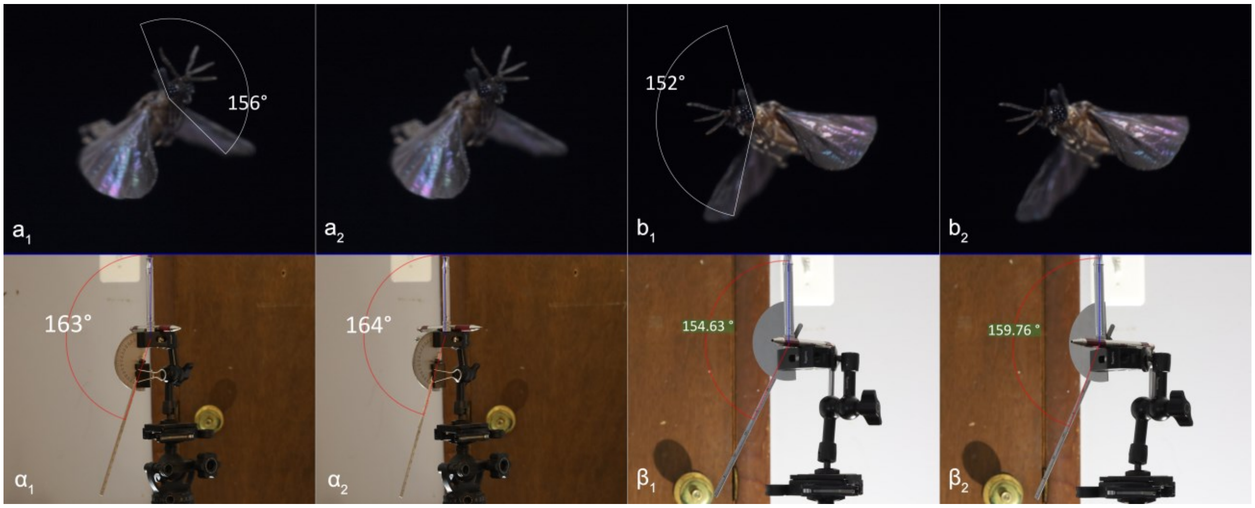


**Fig. S4** Estimated wingstroke amplitudes. Strepsipteran halteres and wings beat in antiphase. Together they can potentially provide a reasonable estimate of the wingstroke amplitude from a still image, provided the halteres are at their apex and the yaw rotation of the body is not more than about 50°. In the upper row, Strepsiptera have been drawn both with the angular measurement between the intersection of the halteres and one of the wings ($a_1$, $b_1$), and, for better landmark identification, without it ($a_2$, $b_2$). In $a_2$ and $b_2$, the leading edge of the (cambered) wing appears as a slight bulge along the periphery of the rearward wing. The lower row are contains images from a series of tests to determine the accuracy of such estimations (see text). In $\alpha_1$ and $\alpha_2$, the pen, which acted as the body axis, is 30° and 40° left of head on; in $\beta_1$ and $\beta_2$, it is first 30°, then 40° right thereof. These angles were displayed because those of the corresponding Strepsiptera above appear to be within this range. Likewise, in $\alpha_1$ and $\alpha_2$, the true angular displacement was 160°, while in $\beta_1$ and $\beta_2$, it was 150°, but the "body" was viewed from slightly below. (a) The halteres are straight above the abdomen, so the wings must be near their nadir. The estimated wingstroke amplitude was measured to be about 156°, indicating that the corrected angle would be about 152°. (b) The estimated wingstroke amplitude was measured to be about 152°, indicating that the corrected angle would be about 147°. Together, the two estimates average out to about 152°. Note, however, that the wings are swept forward somewhat, reducing the apparent discrepancy. Image *(a)* was shot on 17 VIII 2021 @ 5:51 AM; image *(b)* was shot on 25 VIII 2021 @ 4:31 AM. Images $\alpha_1$ and $\alpha_2$ were flipped horizontally, so that the "body" axis would align with that of the insect above it.

---

[1] The wingstroke amplitude could also be estimated when the wings are directly above the abdomen, but there is no guarantee the haltere-stroke has the same (downward) amplitude as the wingstroke.

I ran a series of tests to check if the yaw rotation present in the photos would greatly influence the estimated *T. texana* wingstroke amplitude. I thus used a pen to represent the insect body axis, the mobile straightedge of an adjustable protractor as the leading edge spar of the wing, and a metal rod or screw as one of the halteres. These elements were always eye level with the camera in the α series, but in the β series, a set of measurements was first taken with the pen at eye level, then a second set of measurements was taken when the "insect" was slightly above eye level, and lastly, a third set was taken with the insect slightly above eye level, with the protractor representing its measured wing shifted forward by 30° with respect to perpendicular to the body axis. This was done because the wings are shifted forward in Fig. S4a&b, and as shown in Fig. 3 of the main paper, a 30° produces essentially no bodily friction, although due to the wing camber, that large an offset is an overestimation. Finally, in series α, apparent wingstroke amplitudes were measured for vertical angular wing displacements of 90°, 150°, and 160° (160° shown). In series β, the vertical angular wing displacement was always 150°. The elements were then rotated from 0–60° (α) and from 0–90° (β) in 10° increments.

Findings

For yaw rotations of up to 40°, the largest percent difference from the actual angular wing displacement was +6.5% (estimates were almost always overestimates, and usually within 5%) and −3% (usually within −1.5% across this range). The yaw offset for 30° and 40° are shown in α and β respectively, which appears to encompass the yaw offset of both photographed insects. Displacement from eye-level had little noticeable effect. Yaw offsets ≤ 50° produced results within 7.5% of the actual angular displacement. Thus, the apparent offsets for which one would be likely to make such an estimate all produce surprisingly reasonable results. Yaw offsets of 20° typically produced (small) underestimates. Forward wing displacement ostensibly effectively increased the small yaw displacement by first being slightly in front of flat, ranging to flat, and then becoming a small yaw displacement behind flat, even for much larger overall yaw displacements.

Given these results, the original average estimate of ≈152° is very likely to be within 5% [144°–160°] of the true value, and probably nearer to 3%. Thus, the wingstroke amplitude for *T. texana* is likely somewhat closer to 150° than to 160°, as was found by Pix et al. (1993) for *Xenos vesparum*. Finally, although the halteres nearly touch in Fig. S4b, the angle between their adjacent faces is substantial, whereas they are nearly parallel in Fig. S4a. This may be due to the halteres being pronated at the end of their upstroke, which could signal their evolutionary origin as flight wings.
